# AI for predicting chemical-effect associations at the chemical universe level – deepFPlearn

**DOI:** 10.1101/2021.06.24.449697

**Authors:** Jana Schor, Patrick Scheibe, Matthias Bernt, Wibke Busch, Chih Lai, Jörg Hackermüller

## Abstract

Many chemicals are out there in our environment, and all living species are exposed. However, numerous chemicals pose risks, such as developing severe diseases, if they occur at the wrong time in the wrong place. For the majority of the chemicals, these risks are not known. Chemical risk assessment and subsequent regulation of use require efficient and systematic strategies. Lab-based methods – even if high throughput – are too slow to keep up with the pace of chemical innovation. Existing computational approaches are designed for specific chemical classes or sub-problems but not usable on a large scale. Further, the application range of these approaches is limited by the low amount of available labeled training data.

We present the ready-to-use and stand-alone program `deepFPlearn` that predicts the association between chemical structures and effects on the gene/pathway level using a combined deep learning approach. `deepFPlearn` uses a deep autoencoder for feature reduction before training a deep feedforward neural network to predict the target association. We received good prediction qualities and showed that our feature compression preserves relevant chemical structural information. Using a vast chemical inventory (unlabeled data) as input for the autoencoder did not reduce our prediction quality but allowed capturing a much more comprehensive range of chemical structures. We predict meaningful - experimentally verified-associations of chemicals and effects on unseen data. `deepFPlearn` classifies hundreds of thousands of chemicals in seconds.

We provide `deepFPlearn` as an open-source and flexible tool that can be easily retrained and customized to different application settings at https://github.com/yigbt/deepFPlearn.

**Supplementary information:** Supplementary data are available at bioRxiv online.

## 1 Introduction

### Exposure to a vast amount of chemicals threatens the health of humans and ecosystems

Chemical products are essential for maintaining our standard of living, and chemicals form the building blocks of life. Some chemicals are hazardous upon exposure, and their safety needs to be thoroughly evaluated. The number of chemicals that we are exposed to and the set of chemicals of anthropogenic origin has been rapidly growing from 20 million in 2002 to currently 169 million unique chemicals in The Chemical Abstracts Registry Service Cas [2020]. Many of those chemicals are not relevant for exposure since they are not used in larger quantities. The estimated number of chemicals available on the global market highly varies between 30 000 up to 350 000 Fischer [2017], European Environment Agency [2015], Bond and Garny [2019], Wang et al. [2020]. The number of chemicals detected in human bodies is of similar order of magnitude. Mattingly et al. [2016] compiled more than 50 thousand from scientific texts in the Blood Exposome DB. Rappaport [2016] defined our lifestyle and the change in environmental determinants Lim et al. [2012], Landrigan et al. [2018] rather than genetic factors as the primary cause for many chronic diseases. Contamination with anthropogenic chemicals is of similar concern in the environment Anderson et al. [2006], Busch et al. [2016]. The NORMAN database lists ~3000 chemicals as emerging pollutants across Europe Norman Network [2020]. Chemical exposure was considered a major threat for wildlife populations Köhler and Triebskorn [2013], Hallmann et al. [2014], Desforges et al. [2018]. For example, up to 26% of aquatic species loss may be attributed to exposure to chemical mixtures [European Environment Agency, 2015, p. 245] and Posthuma et al. [2019].

### Risk assessment fails to keep up with the pace of chemical innovation

This enormous chemical exposure and observed hazards require an efficient and effective risk assessment and specific regulation of chemical use. While the EU European Commission [2019] designated a “toxic-free environment” a key priority, the European Environment Agency forecasts that chemical exposure will further increase [European Environment Agency, 2015, pp. 248–249]. For 86% of the ~21 000 chemicals registered under REACH, the European legislation regulating industrial chemicals, the need for suitable regulatory actions still needs to be determined European Chemials Agency [2019]. However, the throughput of regulatory processes is slow compared to the pace of chemical innovation. The evaluation of a chemical of concern takes 7–9 years, during which exposure may continue, under REACH regulation [European Environment Agency, 2015, pp. 248–249]. Also, European Chemicals Agency [2018] found that ~70% of the evaluated registration dossiers are incomplete or not compliant. In summary, traditional approaches to chemical regulation perform well; however, they are too slow to master the number and growth of chemicals on the market. Predictive *in silico* approaches may support this challenge – not necessarily by replacing experimental approaches, but by prioritizing chemicals for further evaluation.

### *In silico* approaches predict toxicity

Different *in silico* approaches exist to predict toxicity. Structural alerts and rule-based models constitute a simple but powerful approach to toxicity prediction and build on using individual chemical substructures as indicators for toxicity Lepailleur et al. [2013]. These methods rely on either humanexpertise-based or data-derived rules and are easy to interpret. However, since no mechanism enforces the completeness of rules, there is a high risk of false-negative predictions Raies and Bajic [2016]. Read-across predicts the unknown toxicity by extrapolation from a set of highly related chemicals with known toxicological properties as an alternative to animal experiments. Naturally, this restricts to the subset of chemicals for which sufficient information of related chemicals is available Vink et al. [2010]. Quantitative Structure Activity Relationships (QSAR) Raies and Bajic [2016] relate molecular descriptors to toxicity or other properties of a chemical Perkins et al. [2003], Cherkasov et al. [2014]. QSARs are built on a set of related chemicals or expert knowledge on several chemicals’ shared mode of action or are derived from a diverse set of chemicals. Descriptors in QSARs include physicochemical properties, different molecular structure representations, or properties thereof, and high throughput screening-derived data on biological activity. A frequently used family of descriptors are molecular fingerprints that record the occurrence of local and regional substructures Lo et al. [2018]. QSAR approaches utilize multivariate statistical models and more recently also machine learning to relate molecular descriptors to toxicity.

### Machine learning in 21st-century toxicology

Toxicology experiences a paradigm shift from relying on apical endpoints in animal models to integrated strategies, combing prediction and high-throughput testing for different endpoints. Screening initiatives, e.g., the Tox21 program Thomas [2018], test thousands of chemicals in hundreds of bioassays to inform on an effect on the molecular processes which are relevant in toxicity. The availability of these data enabled the development of a new variant of QSARs: the machine-learningbased prediction of the association of substances with molecular pathway responses.

The Tox21 Challenge was announced in 2014 to reveal how well independent researchers could predict the interference between chemicals and biochemical pathways given a dataset of chemical structures only. The challenge initiators provided a set of 12 000 chemicals with toxicity effect information for 12 assays along with the task to predict the effects computationally. The winning method was the `DeepTox`[Mayr et al., 2015] pipeline with reported AUC values above 82%. In brief, they normalized the molecular representation of the chemicals, computed a large number of descriptors, and trained deep learning DL models, which they evaluated on the provided test data. Later, the challenge initiators successfully validated these models on withheld data. Further, the authors observed that the models learn simple structures in the lower and more complex structures in the higher layers, as shown for image recognition. In their DL models, hidden neurons represent known molecular substructures - toxicophores, identified manually by experts for decades.

Pu et al. [2019] developed `eToxPred` to quickly estimate the toxicity of extensive collections of low molecular weight organic chemicals. It employs a Restricted Boltzman Machine classifier and a generative probabilistic model to predict a Tox-score. The reported accuracy is 72%.

Sun et al. [2019] used support vector machine and random forest single- and multilabel models to predict toxicity on the Tox21 Challenge data. They used under-sampling to resolve the problem of class imbalance in the data and reported accuracies between 74 to 81%.

Liu et al. [2015] described `TarPred`, a web application for predicting therapeutic and side effect targets of chemicals. It is not available anymore.

The `DeepChem` Project [Ramsundar et al., 2019] is a Python library that provides datasets, functions, and user-contributed tutorials, intending to democratize DL for science in general and chemistry in particular. It is helpful to develop or, as a reference, to compare custom computational approaches in the field.

None of these approaches provides a ready-to-use program for classification or retraining. The main limitation of those (and other) ML approaches in toxicology is the lack of applicability to chemicals outside the training data and the availability of sufficient amounts of training data. A specific challenge is that the descriptors need to be fine-grained enough to capture the particularities of molecular substructures and coarse-grained enough to allow for ML. In particular, the representation of a chemical structure should preserve relevant information which allows for the target association, while it should also summarize structural features to reduce the degrees of freedom of the descriptor space. However, the (high) dimensionality of the features stands in considerable contrast to the (low) amount of available labeled training data.

Here, we present `deepFPlearn` – a ready-to-use DL program that predicts the association of chemical structures and targets on the gene/pathway level. We applied feature reduction via a deep autoencoder (AE) of a simple representation of the chemicals’ structure – the binary fingerprint of moderate size. Subsequently, we predicted the association of the encoded/compressed fingerprint representation with a deep feed-forward neural network (FNN). We overcame the domain extrapolation problem by training the autoencoder on a considerable repertoire of chemical structures and showed that the prediction quality on the subset of labeled training data remained high. Further, we demonstrated that `deepFPlearn` could classify selectively interacting chemicals, which have been experimentally classified recently, with significantly higher confidence than other chemicals.

## 2 Methods and data

### Representation, similarity, and visualization of chemical structures

The molecular structure of chemicals was encoded in binary topological fingerprints – referred to as molecular fingerprint or FP in the following – using Python’s `RDKit` [Landrum, 2006, version 2022.03.1] function `Chem.rdmolops.RDKFingerprint` with minimum path size: 1 bond, maximum path size: 7 bonds, fingerprint size: 2048 bits, number of bits set per hash: 2, minimum fingerprint size: 64 bits, target on-bit density 0.0. These FPs were generated from respective InChI or SMILES strings of the chemicals. All training data files containing chemical structure and target association information were serialized using Python’s pickle module. Uncompressed and compressed features were visualized in 2D space using the UMAP algorithm [McInnes et al., 2018]. To gauge the similarity of two binary FPs *F* and *G* we used the *Tanimoto* similarity - the ratio of intersection and union of set bits:

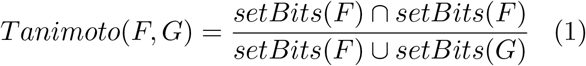

Pearson correlation was used to evaluate the similarity of compressed features. A **k**-means clustering with *k* ∈ [2..7] was applied to the *uncompressed* features. The assigned clusters were translated to color codes in the visualizations of uncompressed and compressed features.

**The deep learning tasks** were implemented using the Python library of the `TensorFlow` framework [Abadi et al., 2016, version 2.6.0] and the `Scikit-learn` framework [Pedregosa FABIANPE-DREGOSA et al., 2011, version 1.0.2].

We used Weights & Biases Biewald [2020] for experiment tracking and hyperparameter tuning (sweep). Sweeped parameters included activation function for the hidden layers, optimizer, learning rate, learning rate decay, batch size, and dropout. The supplement provides all details about the hyperparameter tuning procedure and the final selected training parameter values in section *Hyperparameter tuning*.

We applied a stratified train-test splitting to keep the same distribution of class labels in both, the training and the test data. For training the FNN models, we applied a stratified *k*-fold (default: *k* = 5) cross-validation. We enabled early stopping and fallback mechanisms and monitored validation loss (*val-loss*). The training stopped early if *val-loss* did not improve by *min*Δ = 0.0001 for a certain number of epochs (*patience*_AE_ = 5, *patience*_FNN_ = 20). The model’s weights were restored to the respective checkpoint model. `deepFPlearn` saves the model weights for each fold and the model that performed best across all *k* folds for subsequent prediction and further application. Training histories in terms of the values for loss, binary accuracy, area under the reciever-operator curve (AUC-ROC), precision, recall, and F1 score were logged in .CSV format for each training epoch’s training and validation data. To find the optimal classification threshold we used Matthews Correlation Coefficient (MCC) as one of the unbiased evaluation metrics for imbalanced classification. MCC was calculated with an increasing threshold from 0 to 1 on the predicted values of the validation data (and their true values). Then, the threshold with maximum MCC was selected as the *tuned* classification threshold for each model individually. See Figure **??** C for an example.

**The deep regularized autoencoder** has a symmetric shape with one-dimensional input and output layers of the size of the input fingerprint *L_FP_*. The number of hidden layers *N_H_* and their sizes *S_i_, i* ∈ [1..*N_H_*] depend on *L_FP_* and the desired size of the latent space *L_z_* – see Table 1 for the applied sizes of the hidden layers.

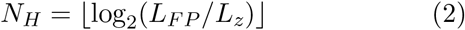

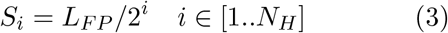

**Table 1:** Number and sizes of hidden layers for each trained neural network. NN – neural network; AE – autoencoder; FNN – feed-forward NN

| NN | Input | Input size | hidden layers |
| --- | --- | --- | --- |
| AE | FP | $L_{FP} = 2048$ | 1024, 512, 256, 512, 1024 |
| FNN | FP | $L_{FP} = 2048$ | 1024, 512, 256, 128 |
| FNN | compressed FP | $L_z = 256$ | 128, 64, 32 |

The SELU activation function and *lecun-normal* weight initialization were used in hidden layers, and the Sigmoid activation function for the output layer. The model was compiled with binary cross-entropy as loss function and Adam optimizer.

**A deep neural network** was constructed as a sequential feed-forward neural network for the classification task. The dimensions of the stacked layers depend on the mode of action of `deepFPlearn`. If feature compression via the AE is enabled, the FNN is used subsequently. Then, the size of the input layer of the FNN matches the length of the latent space *L_z_* vectors. Otherwise, the size of the input layer matches the length of the molecular fingerprint *L_FP_*. The hidden layers were of decreasing sizes, followed by an output layer of size 1. The number of hidden layers 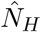 and their sizes 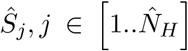 depend on the provided input size *L_input_* (which is either *L_FP_* or *L_z_*). The last four layers with only a few neurons (e.g. less than 32 in the example case of *L_FP_* = 2048) were not included. See Table 1 for the applied sizes of the hidden layers.

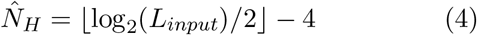

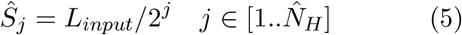

Dense layers were used with the SELU activation function, *lecun-normal* weight initialization, and *AlphaDropout*. The Sigmoid activation function was used for the output layer. All hidden layers were followed by a dropout layer. To reflect a potential imbalance in the training data, we introduced an initial bias of log(*P/N*) (with *P* equal to the number of 1-values and *N* to the number of 0-values in the target vector) to the output layer. The FNN model was compiled using the Adam optimizer and binary cross entropy as loss function

**Different datasets** were collected from the literature and public databases. A manually curated dataset *S* was downloaded from the supplemental material of Sun et al. [2019]. It contained chemical-target associations for 7248 chemicals and 6 gene targets that are involved in endocrine disruption (ED) in humans (*androgen receptor* (AR), *estrogen receptor* (ER), *glucochorticoid receptor* (GR), *thyroid receptor* (TR), *PPARg*, and *Aromatase*). See supplemental Fig. S1 for an overview of this data’s size and class distributions. Initially, this data had been retrieved from bioassay data of the Tox21 program [Thomas, 2018], and carefully transformed to binary associations by Sun et al. [2019]: Associations were considered as not available (NA) if no bioassay data was available, and as 1 or 0, if an association between chemical and gene target had been confirmed in a bioassay or not, respectively. See Sun et al. [2019] for details. The dataset *S* was extended by the artificial target ED that combines all existing target associations with a logical OR operation. Chemicals in the *S* dataset were identified by their SMILES string.

Further, a dataset *D* was generated from the 719 996 chemicals listed in the CompTox Chemistry Dashboard [Williams et al., 2017, accessed on 2020/07/13]. Chemicals in the *D* dataset were identified by their InChI identifiers.

For benchmarking, we downloaded two datasets from MoleculeNet Wu et al. [2017], a database of benchmarking datasets for classification problems in molecular ML. First, we selected the Tox21 Challenge dataset - *Tox*21, which associates chemicals and gene targets. Second, we used the Side Effect Resource (*SIDER*) database that associates drugs with grouped adverse drug reactions. These datasets contained 7831 and 1427 compounds, and 12 and 27 targets, respectively, and comprised binarized associations between those compounds and the targets. We followed the recommended metric and splitting patterns Wu et al. [2017] to generate training data from these datasets and selected targets with a 1-0-ratio of at least 0.2 and a minimal number of 200 samples in the positive class for training.

See Table 2 for an overview which of those datasets were used in which training or prediction case.

**Table 2:** Overview of the usage of the different datasets for training or prediction.

| case | train AE | use AE | train FNN | predict |
| --- | --- | --- | --- | --- |
| 1 | *S* | - | - | - |
| 2 | *D* | - | - | - |
| 3 | - | - | *S* | - |
| 4 | - | *S* | *S* | - |
| 5 | - | *D* | *S* | - |
| 6 | - | *D* | *SIDER* | - |
| 7 | - | *D* | *Tox21* | - |
| 8 | - | *D* | - | *D* |

### Implementation

`deepFPlearn` was implemented as a Python (version 3.9.12) package with three different usage-modes. First, *convert* imports the dataset (for training or prediction) and calculates molecular fingerprints for all structures from their respective SMILES or InChi representation. A data frame combines the original representation, the calculated fingerprint, and all targets. It is then serialized to disc as a Pickle file to accelerate the data import for subsequent sessions. Importantly, `deepFPlearn` assumes that SMILES have been canonicalized and cleaned. We recommend to either use ChemAxon’s chemical structure representation toolkit^1^ or a chemical structure curation pipeline relying on RDKit [Bento et al., 2020]. The second mode is *training*. The neural networks can easily be (re-)trained with any dataset that associates chemical structures with an effect. All necessary information is logged during the training to validate and evaluate the trained models. The third mode is to *predict* the association of a provided list of chemicals with an effect using the trained models.

The user can adjust all neural network settings and the mode of action in a JSON configuration file.

Dependencies to external libraries and software are managed using a platform-independent conda^2^ environment, which we provide in the code repository. A singularity container^3^ was set up that encapsulates the whole project at the state of publication for usage and reproducibility. It includes the required resources, source code, compiled package, and test data.

## 3 Results

We developed the stand-alone, ready-to-use DL approach `deepFPlearn` to associate chemicals with gene/pathway level targets. We further evaluated the potential of feature compression to increase the applicability to substances beyond the limited amount of available training data.

Our workflow combined a pre-training strategy via a deep autoencoder to reduce the feature space and to generate a universal encoding of binary fingerpints, followed by a classification step using a deep feed-forward neural network, see Figure **??**.

For the FNNs, we employed 5-fold crossvalidation to show that the selection of the traintest-split has no significant impact on the model performance. In particular, the standard deviation of the ROC-AUC values (calculated on the validation data) was ~1%, see supplement Fig. S2. Therefore we used a single stratified train-test-split to finetune and train our models.

### Feature compression comprehensively reduced trainable parameters while keeping comparable classification performance

We applied different training setups: First, feature compression was disabled (*no* AE, Figure **??** from A to C), and FNN training used the full-length molecular fingerprints. The ratio between positive (1) and negative (0) associations differed substantially between the individual targets, see supplement Fig. S1. We introduced an initial bias to the output layer of the FNN to reflect that imbalance and selected AR, ER, and the artificial target ED as subsets with an acceptable imbalance to train individual FNN models. Due to the fingerprint size of 2048, the respective hidden layer sizes of the FNN were 1024, 512, 256 and 128 resulting in about 2.8 × 10^6^ trainable parameters. The training stopped early before ~ 100 epochs. Binary accuracy values of 0.85, 0.83 and 0.78 and ROC-AUC values of 0.81, 0.83 and 0.81 were reached for AR, ER, and ED, respectively. See Figure **??** A (top panels) for the training histories, and Figure **??** B (lightgrey bars) for the values of precision, recall, F1 scores and further metrics that describe the performance of our FNN models. See Figure **??** A for ROC and precision-recall curves of the AR target for the classification without AE, and supplement Fig. S 3–5 for confusion matrices, ROC and precision-recall curves of all three targets.

Second, we applied feature reduction before the classification by training an AE with a latent space size of *L_z_* = 256. This reduced the respective hidden layer sizes of the FNN to 128, 64 and 32, resulting in only 43.3 × 10^3^ trainable parameters, which is 1.55% of the uncompressed case above. We trained both a *specific* AE using the (small) *S* dataset and a *generic* AE using the (large) *D* dataset. See Figure **??** from A over B to C. The training of the specific autoencoder stopped early at 28 epochs which is due to the small number of training samples. The validation loss reached a value of 0.026. The generic autoencoder trained for around 320 epochs and stopped at a validation loss of 0.159. See Figure **??** for the training histories and a UMAP visualization of the high-dimensional uncompressed feature space and the low-dimensional latent space of dataset *S*. Coloring compounds from the uncompressed and compressed space with labels calculated on the uncompressed feature space yielded similar cluster associations in the UMAP. Therefore, the AE preserves relevant (structural) information during feature compression.

Subsequently, we trained the FNNs and used the latent space representation as input. The training stopped early before ~400 epochs. Binary accuracy values of 0.85, 0.80 and 0.77 and ROC-AUC values of 0.81, 0.81 and 0.78 were reached for AR, ER, and ED, respectively, when the specific AE was used to encode the fingerprints. We observed no significant discrepancy in these values when using the generic AE. In particular, we reached values of 0.85, 0.80 and 0.74 for binary accuracy, and 0.80, 0.79 and 0.76 for ROC-AUC values. Therefore, when the input features are compressed with the generic AE, the FNNs may be applied to a much more comprehensive range of molecular structures without compromising on the predictive power. See Figure **??** A (middle and lower panels) for the training histories, and Figure **??** B (medium and dark grey bars) for the values of precision, recall, F1 score and further metrics that describe the performance of our FNN models that were trained with compressed fingerprints. See Figure **??** B for ROC and precision-recall curves of the AR target for the classification with the generic AE, and supplementary Fig. 3 – 5 for confusion matrices, ROC and precision-recall curves for all three targets.

### Benchmarking confirmed our strategy

We compared the results of our strategy against the results of Sun et al. [2019], the publication from which we extracted our FNN training data, the introduced approaches `eToxPred` Pu et al. [2019] and `DeepTox` Mayr et al. [2015], and the results reported by MoleculeNet Wu et al. [2017]. Sun et al. [2019] reported balanced accuracy values in their results and we reached the same range between 74 to 81% on the same data. Pu et al. [2019], Mayr et al. [2015] and Wu et al. [2017] reported ROC-AUC values of 72, 82 and 83%, respectively, on the Tox21 data of MoleculeNet while our models achieved ROC-AUC values of 88%. For the SIDER dataset Wu et al. [2017] reported 67% ROC-AUC values while we reached 84%. For the MoleculeNet datasets, we also observed only a slight drop in performance when using the generic AE. In summary, our models perform either in the same range as existing approaches or better, which is satisfying compared to the increased applicability of our strategy.

### deepFPlearn is ready to be applied to huge datasets

We used `deepFPlearn` with generic feature compression and selected the trained models for AR, ER and ED to predict associations of the ~ 700*k* chemicals from dataset *D*. For most of those compounds, the probability of acting as endocrine disruptors was not known. `deepFPlearn` predicted ~60k with high prediction probability *P* > 0.85. From the ED predictions of dataset *D*, we investigated the top 200 and bottom 200 (ranked by prediction probability) and empirically investigated their biological feasibility. We found compounds among the top 200 like Estriol, 17alpha-Ethinylestradiol, 17beta-Ethinylestradiol, Mestranol, Prednisolone Dexamethasone, Betamethasone, and respective derivates. These chemicals are well known to interact with the human estrogen receptors and pathways or with the glucocorticoid pathway. Interestingly, Escher et al. [2020] also identified some of those to interact selectively with AR in the cell assay screenings. Also, the top 200 list contains the chemicals Ezlopitant dihydrate dihydrochloride, 5-Bromo-2, 2-diethyl-5-nitro-1,3-dioxane, or Schinifoline, a metabolite of the Japanese Pepper plant Zanthoxylum schinifolium. To our knowledge, those substances have not been tested in bioassays so far. In the bottom 200 predictions (*P* < 0.01) we found derivates of carbamic, acetic, and amino acids. Those chemicals have never been discussed in the context of steroid hormone related ED as far as we know.

Recently, Escher et al. [2020] categorized a selection of 355 out of 7968 investigated chemicals and their activity with the ED receptors AR and ER as *selective* (41), *specific* and *unspecific* (314, summarized as *other*) binders.

We predicted the associations for the subset of 339 chemicals that have not been part of our training data with and without generic feature compresssion. The models for ER and ED that were trained on the compressed fingerprints captured substantially more of the selective compounds with higher prediction probability than the models that used the uncompressed fingerprints. However, this was not true for the AR model. See Figure **??** B and C for probability distributions and counts of the ED model and supplement Fig. S6 for the comparison of all three models.

## 4 Discussion

There is a great need for systematic prediction of chemical-effect associations in toxicology. They are required to prioritize chemicals for experimental screening, a smart selection of chemicals for monitoring, and the design of novel chemicals. Several approaches and implementations exist that partially address these challenges. However, no tools for large-scale application are available, and the option for retraining with additional data sets is absent. While MoleculeNet [Wu et al., 2017] and Deepchem [Ramsundar et al., 2019] provide capable frameworks for developing learning applications on chemicals, readily applicable tools, e.g., for predicting ED, are missing.

With `deepFPlearn` we present an application to investigate sets of chemicals for their potential associations to gene targets involved in ED. It is a DL approach with the possibility of training custom models to predict different associations of interest.

The small number of labeled training data is in contrast to the high number of features necessary to describe a chemical’s molecular structure. Also, the natural interaction of chemicals and biomolecules is biased towards “no interaction” (label of 0) such that the data suffer from a substantial imbalance between 1 and 0 labels. Assessing the association of chemicals and biomolecules requires measuring a range of concentrations per substance and assay and thus poses a substantial effort even with high-throughput technologies. Since the number of substances with measured associations is small compared to the universe of chemicals, there is a lack of labeled training data. Due to the high speed at which new chemicals are developed, this situation will not change in the foreseeable future. To make things worse, many positive associations (label of 1) are potentially wrong due to mistakes during screening result interpretation. Examples are unclear effect thresholds, high variability in the experimental designs, and limitations in the statistics of modeling the observed effect. The imbalance of the training data together with a large number of parameters can easily lead to overfitting. However, our strategy to initialize the output layer of the FNN with the correct bias to reflect the imbalanced prevented overfitting also for the FNN.

We reduced the discrepancy between large descriptor size and the limited training data by compressing features with a deep autoencoder. Further, this reduced the large number of trainable parameters to 1.55% of the networks that do not use an AE. Using a large repertoire of chemicals for training the AE further improved the domain extrapolation without reducing the predictive power of the subsequent classification. We tested different training situations, i) without feature compression, ii) feature compression with a subset of chemicals (specific AE), and iii) feature compression with a large set of chemicals (generic AE). We reached good training performances with ROC-AUC values above 80%, with satisfying sensitivity up to 75%, and specificity up to 97%.

Using the benchmark datasets from MoleculeNet, and reported binary accuracies and ROC-AUC values from other approaches that used the same data sets we showed that `deepFPlearn` performed comparably or better. However, those methods also demand significant adjustments to the training data to cope with imbalance. We found that our predictions with the generic AE captured more of the compounds that have been experimentally analyzed and classified by Escher et al. [2020] than the models trained on the uncompressed fingerprints, which verifies our assumption on predicting unseen data.

`deepFPlearn` allows for selecting different usage modes depending on the classification problem: If the compounds to be classified are expected to reside within the domain of the training data the FNN without AE provides superior classification performance. However, given the overall comparable accuracy of `deepFPlearn` when pre-training on a large data set, we consider this the more robust, computationally efficient, and generally more applicable approach in particular for large, heterogeneous, and imbalanced data.

The quality of our predictions is also high on the large CompTox dataset. Among the top 1– associated predictions were chemicals that are well known to interact with human estrogen or the glucocorticoid receptor or related pathways. Likewise, among the respective top 0–associated predictions were chemicals that have never been discussed to be involved in ED, which further enhances the confidence in our models.

Our high values for specificity also suggest an application of `deepFPlearn` to predict secondary effects in drug design.

The `deepFPlearn` results on the chemicals experimentally classified as selective and unspecific also confirmed our prediction quality. Although a relatively broad distribution of prediction probabilities for selective binders suggests that there is still room for methodological improvement, many of the chemicals predicted with a very high probability are indeed selective binders.

We suggest a more detailed investigation of the predicted associations and experimental validation in upcoming studies to confirm or decline effects in endocrine disruption.

## 5 Conclusion

With `deepFPlearn` we model the associations between chemical structures and effects on the gene/pathway level with a deep learning approach.

In contrast to existing approaches and implementations, `deepFPlearn` is a ready-to-use tool. It comes as a stand-alone Python software package and (additionally) wrapped in a Singularity Container to overcome the dependency on the operating system and required software. `deepFPlearn` can capture a much more comprehensive range of substances than those contained in the training data of the classification network. It can be applied to classify hundreds of thousands of chemicals in seconds. Moreover, with its different application modes, we provide the flexibility to train custom models with any meaningful dataset that associates chemicals with an effect. `deepFPlearn` substantially contributes to the systematic *in silico* investigation of chemicals, even for data-driven hypothesis generation on novel substance-effect associations. With `deepFPlearn` we can cope with the large, constantly, and rapidly growing chemical universe and support prioritization of chemicals for experimental testing, assist in the smart selection of chemicals for monitoring and contribute to the sustainable design of the future chemicals.

## Supporting information

Specific training data set S

CompTox data set D

## 6 Key Points

- All living species are exposed to a vast amount (and mixtures) of chemicals; many pose risks; this risk is not known for the majority.
- To support the lab-based risk assessment and subsequent regulation of use, prioritize chemicals for experimental design and hypothesis generation, efficient and systematic tools that evaluate the chemical-effect association on a large scale are required but not available so far.
- We present the ready-to-use deep learning application `deepFPlearn` that predicts the association between the chemical’s molecular structure and the observed effect on the gene/pathway level.
- We solved the discrepancy between large feature space describing the molecular structure and the low amount of labeled training data with a pre-training strategy for feature compression on the chemical inventory.
- We confirmed the good performance and high prediction quality of `deepFPlearn` with benchmarking and experimentally validated datasets.

## 7 Availability

The source code is available in a git repository at `github`: https://github.com/yigbt/deepFPlearn under the terms of the UFZ license, which is based on GNU General Public License as published by the Free Software Foundation version 3 or later. We refer to this repository for installation and usage instructions. For ease of use we also provide Docker and Singularity containers, which is accessible via this repository.

## 8 Competing interests

The authors declare that they have no competing interests.

## 9 Author contributions statement

JS and JH planned the study; JS and CL defined the neural network architectures; JS preprocessed all data; JS, PS implemented the software package and analyzed the results; MB and PS built the singularity container and all github actions; All authors wrote the manuscript. All authors read and approved the final manuscript.

## 10 Acknowledgments

The authors are grateful to Martin Krauss for helpful discussions. This work was supported in part by the Helmholtz.AI project XAI-graph and by the CEFIC Long Range Initiative through funding the project C5 - XomeTox.

---

**Figure 1:**
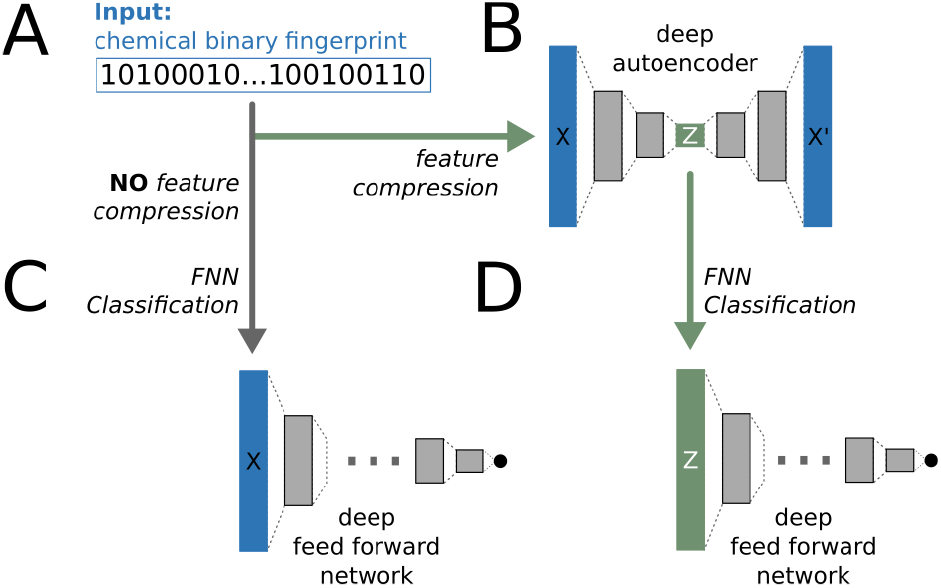
The `deepFPlearn` workflow. A) The molecular fingerprints serve as input for the neural networks. B) A deep autoencoder (AE) is used to compress the fingerprints. C) A deep feed forward network (FNN) is used for direct classification of the input. D) A deep feed forward network (FNN) is used for classification of the compressed input. Sizes of layers, activation and loss functions are different for each network and depend on the input size, see methods section.

**Figure 2:**
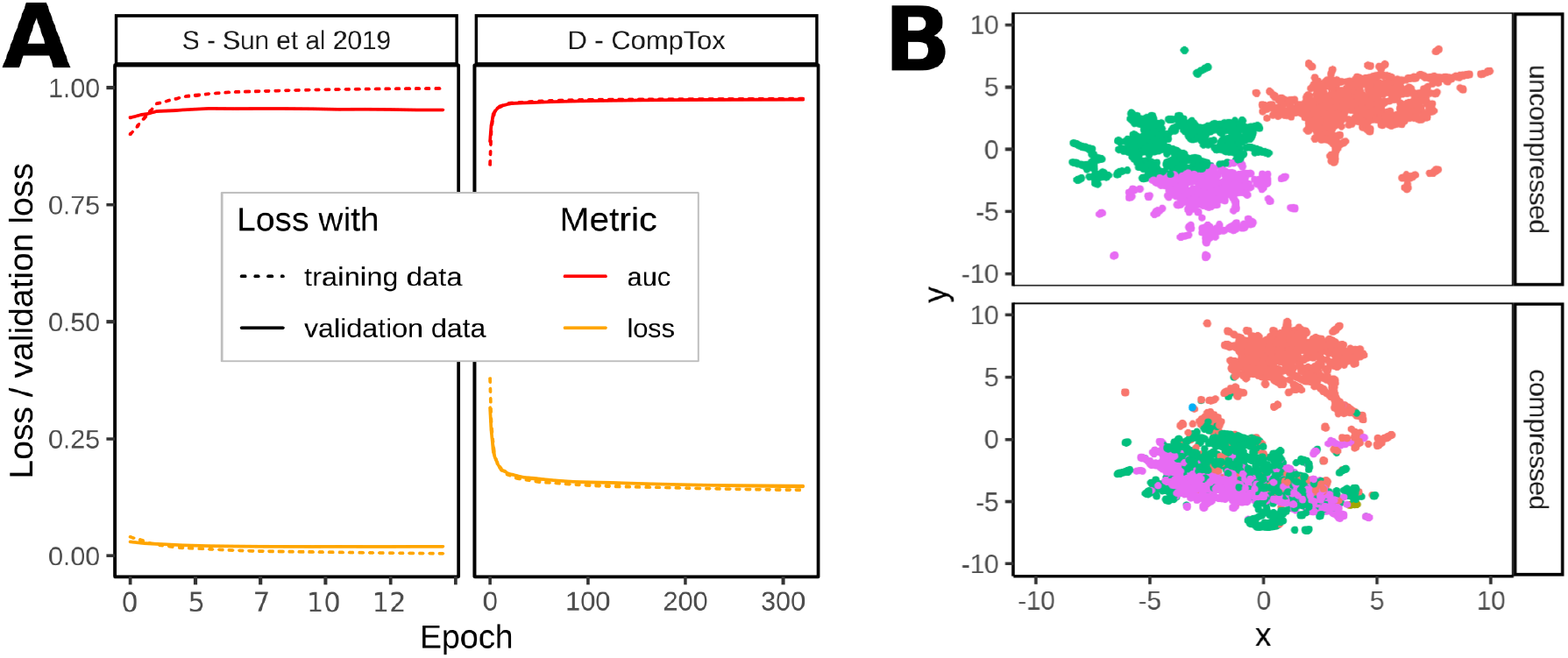
A) ROC-AUC and loss values during training (calculated on the training and validation data after each epoch) of the *specific* (*S* - Sun et al. 2019) and the *generic* (*D* - CompTox) autoencoder. The training stopped early at 28 epochs for the specific AE - due to the small number of available training samples and reached a validation loss of 0.026. The training of the generic AE stopped at ~320 epochs reaching a validation loss of 0.159. B) UMAP visualizations of uncompressed and compressed representations of all compounds from *S* dataset; color indicates cluster assignment of a *k*-means clustering with *k* = 4 on the uncompressed features.

**Figure 3:**
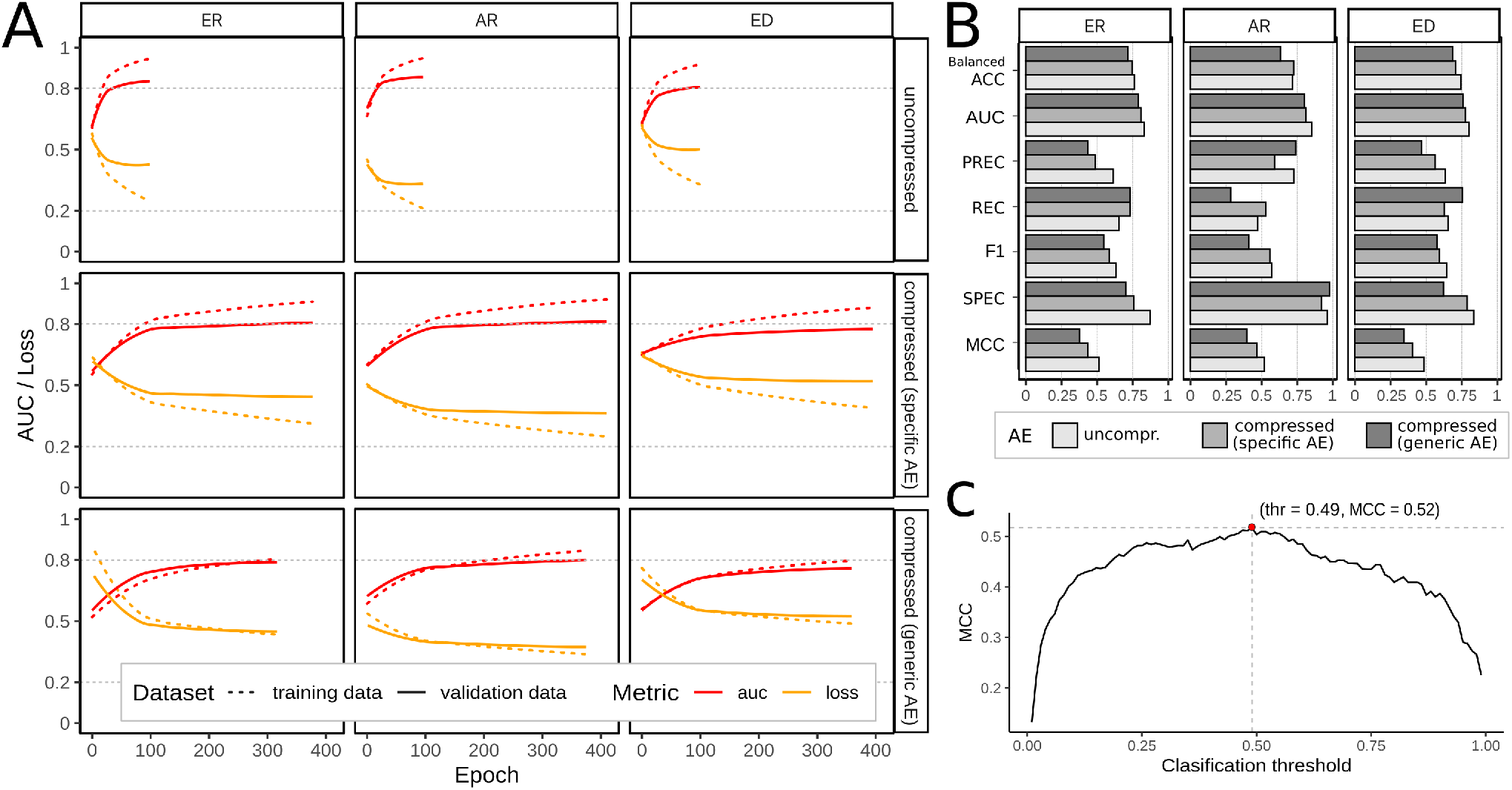
A) Training histories of the feed forward neural networks stratified by the selected targets/models for androgen (AR) and estrogen (ER) receptors, and endocrine disruption (ED), and the degree of feature compression (uncompressed, specific AE, and generic AE); shown metrics are ROC-AUC (red), loss (orange) calculated on the training (dotted) and validation data (solid) during training. B) Comparison of the values of balanced accuracy (Balanced ACC), area under the receiver-operator curve (AUC), precision (PREC), recall (REC), F1 score (F1), specificity (SPEC), and Matthews correlation coefficient (MCC) of the individual models using no (lightgrey), the specific (medium grey) and the generic AE (dark grey). C) MCC was calculated for increasing thresholds from 0 to 1 on the predicted validation data. The threshold with maximum MCC was selected as the individual classification threshold for each model. Example generated for model: AR, uncompressed input.

**Figure 4:**
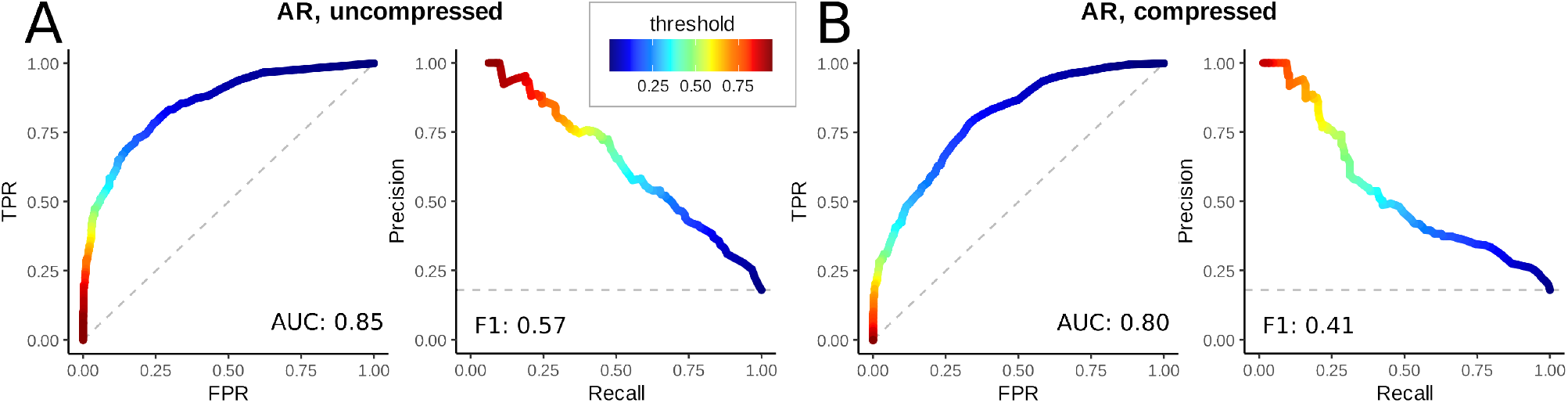
Receiver-operator (left of both panels) and precision-recall (right of both panels) curves of the AR target without using feature compression (A), and with generic feature compression (B). The color indicates the value of the respective classification threshold.

**Figure 5:**
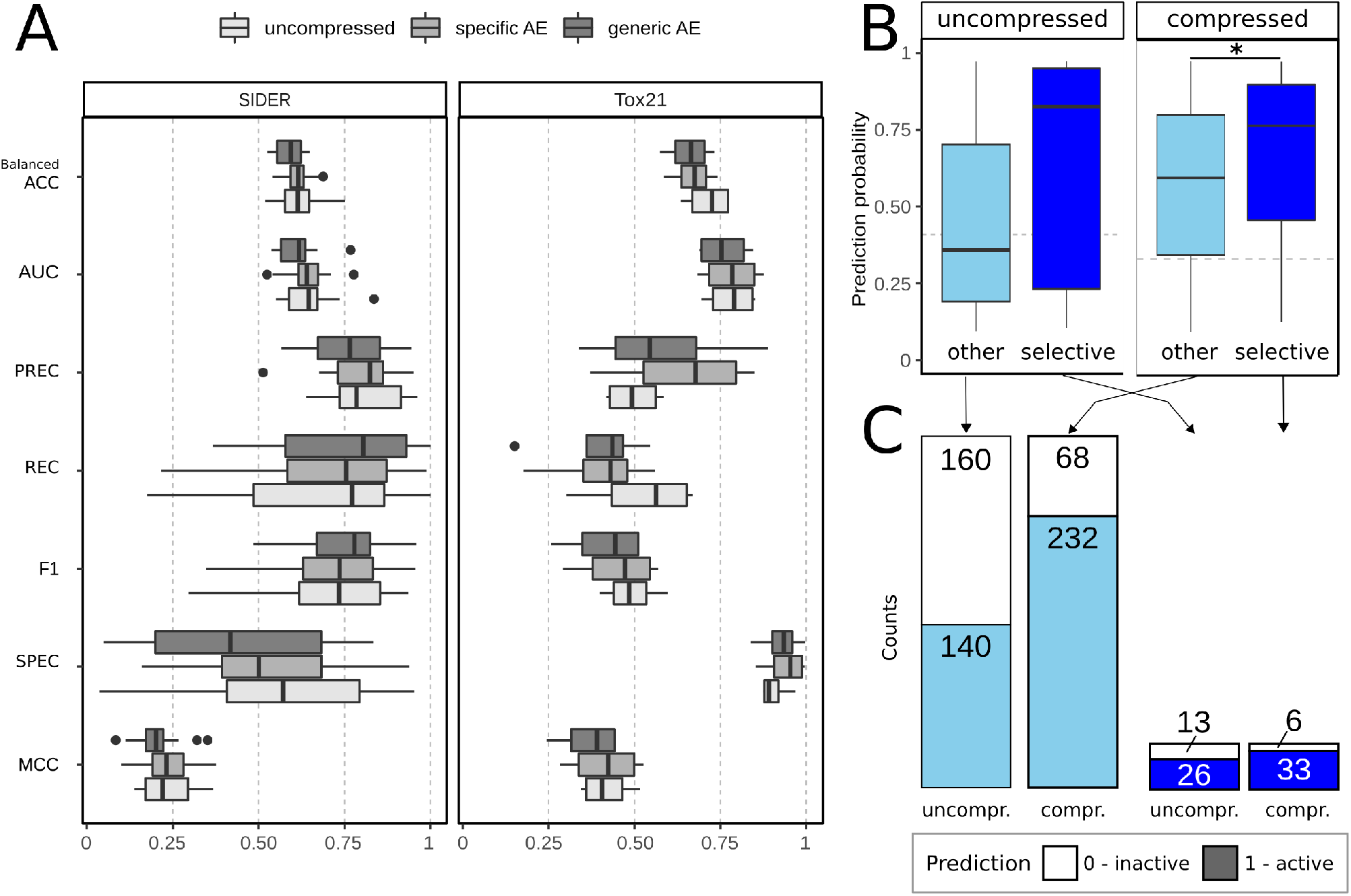
A) Values for all metrics calculated on the validation data for the benchmarking data sets SIDER and Tox21 summarized across all targets: balanced accuracy (Balanced ACC), area und the receiver operator curve (AUC), precision (PREC), recall (REC), F1 score (F1), specificity (SPEC), and Matthews correlation coefficient (MCC) of the individual models using no (light grey), the specific (medium grey) and the generic AE (dark grey). B) `deepFPlearn` prediction probabilities using the ED model with generic AE on the compounds that have been experimentally measured for quantified target association and respectively differentiated into selective and non-specifically acting compounds by Escher et al. [2020]. Probability distributions are compared using the Kolmogorow-Smirnow test, and the significance levels for rejecting the null hypotheses that both distributions are similar was * for p-values below 0.05. C) Comparison of the counts of predicted 1 (active) and 0 (inactive) labels for the same compounds as described in Figure B shown for the ED model.

## Footnotes

1 https://chemaxon.com/products/chemical-structure-representation-toolkit

2 https://www.anaconda.com

3 https://sylabs.io/

## Notes

https://github.com/yigbt/deepFPlearn

### Competing Interest Statement

The authors have declared no competing interest.

### Summary of Updates

We have revised our hyperparameter tuning and integrated initial bias to the output layer of our FNNs to reflect the data imbalance. This improved our model performances and the subsequent application range of deepFPlearn.

## References

M. Abadi, P. Barham, J. Chen, Z. Chen, A. Davis, J. Dean, M. Devin, S. Ghemawat, G. Irving, M. Isard, M. Kudlur, J. Levenberg, R. Monga, S. Moore, D. G. Murray, B. Steiner, P. Tucker, V. Vasudevan, P. Warden, M. Wicke, Y. Yu, and X. Zheng. TensorFlow: A system for large-scale machine learning. In Proceedings of the 12th USENIX Symposium on Operating Systems Design and Implementation, OSDI2016, pages 265–283, 2016. ISBN 9781931971331. URL https://tensorflow.org.

M. G. Anderson, J. Mcdonnell, C. Ximing, S. A. Cline, W. W. Balance, J. Rockstrom, G. C. Daily, P. R. Ehrlich, C. A. Reidy, M. Dynesius, C. Revenga, L. Dams, E. Reichel, M. A. Global, P. Green, J. Salisbury, R. B. Lammers, S. Kanae, T. Oki, A. Y. Hoekstra, P. Eichert, K. C. Abbaspour, A. B. Zehnder, A. Kitoh, M. Hosaka, K. A. Dunne, A. V. Vecchia, C. Valeo, K. Heal, I. Science, M. Bengtsson, Y. Agata, H. Kim, and E. Science. The Challenge of Micropollutants in Aquatic Systems. Science, 313(August):1072–1077, 2006.

A. P. Bento, A. Hersey, E. Félix, G. Landrum, A. Gaulton, F. Atkinson, L. J. Bellis, M. D. Veij, and A. R. Leach. An open source chemical structure curation pipeline using rdkit. Journal of Cheminformatics, 12, 9 2020. ISSN 17582946. doi: 10.1186/s13321-020-00456-1.

L. Biewald. Experiment tracking with weights and biases, 2020. URL https://www.wandb.com/. Software available from wandb.com.

G. G. Bond and V. Garny. Inventory and evaluation of publicly available sources of information on hazards and risks of industrial chemicals. Toxicology and Industrial Health, 35(11-12):738–751, nov 2019. ISSN 0748-2337. doi: 10.1177/0748233719893198.

W. Busch, S. Schmidt, R. Kühne, T. Schulze, M. Krauss, and R. Altenburger. Micropollutants in European rivers: A mode of action survey to support the development of effect-based tools for water monitoring. Environmental Toxicology and Chemistry, 35(8):1887–1899, 2016. ISSN 15528618. doi: 10.1002/etc.3460.

Cas. No Title. https://www.cas.org/support/documentation/chemical-substances, 2020.

A. Cherkasov, E. N. Muratov, D. Fourches, A. Varnek, I. I. Baskin, M. Cronin, J. Dearden, P. Gramatica, Y. C. Martin, R. Todeschini, V. Consonni, V. E. Kuz’Min, R. Cramer, R. Benigni, C. Yang, J. Rathman, L. Terfloth, J. Gasteiger, A. Richard, and A. Tropsha. QSAR modeling: Where have you been? Where are you going to? Journal of Medicinal Chemistry, 57(12):4977–5010, 2014. ISSN 15204804. doi: 10.1021/jm4004285.

J. P. Desforges, A. Hall, B. McConnell, A. Rosing-Asvid, J. L. Barber, A. Brownlow, S. De Guise, I. Eulaers, P. D. Jepson, R. J. Letcher, M. Levin, P. S. Ross, F. Samarra, G. Víkingson, C. Sonne, and R. Dietz. Predicting global killer whale population collapse from PCB pollution. Science, 361(6409):1373–1376, 2018. ISSN 10959203. doi: 10.1126/science.aat1953.

B. I. Escher, L. Henneberger, M. König, R. Schlichting, and F. C. Fischer. Cytotoxicity burst? Differentiating specific from nonspecific effects in tox21 in vitro reporter gene assays. Environmental Health Perspectives, 128(7):1–10, jul 2020. ISSN 15529924. doi: 10.1289/EHP6664. URL https://ehp.niehs.nih.gov/doi/10.1289/EHP6664.

European Chemials Agency. No Title. https://echa.europa.eu/de/universe-of-registered-substances, 2019.

European Chemicals Agency. Evaluation under REACH: progress report 2017 — 10 years of experience. Technical report, European Chemicals Agency, Helsinki, 2018. URL https://echa.europa.eu/documents/10162/13628/evaluation_under_reach_progress_en.pdf.

European Commission. COMMUNICATION FROM THE COMMISSION TO THE EUROPEAN PARLIAMENT, THE EUROPEAN COUNCIL, THE COUNCIL, THE EUROPEAN ECONOMIC AND SOCIAL COMMITTEE AND THE COMMITTEE OF THE REGIONS. The European Green Deal. Technical Report COM(2019) 640 final, European Commission, 2019. URL https://ec.europa.eu/info/sites/info/files/european-green-deal-communication_en.pdf.

European Environment Agency. State and Outlook 2015 the European Environment. Technical report, European Environment Agency, 2015.

S. Fischer. KEMI Market List (Version NORMANSLE-S17.0.1.4). http://doi.org/10.5281/zenodo.3959394, 2017.

C. A. Hallmann, R. P. B. Foppen, C. A. M. Van Turnhout, H. De Kroon, and E. Jongejans. Declines in insectivorous birds are associated with high neonicotinoid concentrations. Nature, 511(7509):341–343, 2014. ISSN 14764687. doi: 10.1038/nature13531.

H. R. Köhler and R. Triebskorn. Wildlife ecotoxicology of pesticides: Can we track effects to the population level and beyond? Science, 341(6147):759–765, 2013. ISSN 10959203. doi: 10.1126/science.1237591.

P. J. Landrigan, R. Fuller, N. J. R. Acosta, O. Adeyi, R. Arnold, N. N. Basu, A. B. Baldé, R. Bertollini, S. Bose-O’Reilly, J. I. Boufford, P. N. Breysse, T. Chiles, C. Mahidol, A. M. Coll-Seck, M. L. Cropper, J. Fobil, V. Fuster, M. Greenstone, A. Haines, D. Hanrahan, D. Hunter, M. Khare, A. Krupnick, B. Lanphear, B. Lohani, K. Martin, K. V. Mathiasen, M. A. McTeer, C. J. L. Murray, J. D. Ndahimananjara, F. Perera, J. Potočnik, A. S. Preker, J. Ramesh, J. Rockström, C. Salinas, L. D. Samson, K. Sandilya, P. D. Sly, K. R. Smith, A. Steiner, R. B. Stewart, W. A. Suk, O. C. P. van Schayck, G. N. Yadama, K. Yumkella, and M. Zhong. The Lancet Commission on pollution and health. Lancet (London, England), 391(10119):462–512, 2018. ISSN 1474-547X. doi: 10.1016/S0140-6736(17)32345-0. URL http://www.ncbi.nlm.nih.gov/pubmed/29056410.

G. Landrum. RDKit: Open-source Cheminformatics, 2006. ISSN 00028282.

A. Lepailleur, G. Poezevara, and R. Bureau. Automated detection of structural alerts (chemical fragments) in (eco)toxicology. Computational and Structural Biotechnology Journal, 5(6): e201302013, 2013. ISSN 20010370. doi: 10.5936/csbj.201302013. URL http://dx.doi.org/10.5936/csbj.201302013.

S. S. Lim, T. Vos, A. D. Flaxman, G. Danaei, K. Shibuya, H. Adair-Rohani, M. Amann, H. R. Anderson, K. G. Andrews, M. Aryee, C. Atkinson, L. J. Bacchus, A. N. Bahalim, K. Balakrishnan, J. Balmes, S. Barker-Collo, A. Baxter, M. L. Bell, J. D. Blore, F. Blyth, C. Bonner, G. Borges, R. Bourne, M. Boussinesq, M. Brauer, P. Brooks, N. G. Bruce, B. Brunekreef, C. Bryan-Hancock, C. Bucello, R. Buchbinder, F. Bull, R. T. Burnett, T. E. Byers, B. Calabria, J. Carapetis, E. Carnahan, Z. Chafe, F. Charlson, H. Chen, J. S. Chen, A. T.-A. Cheng, J. C. Child, A. Cohen, K. E. Colson, B. C. Cowie, S. Darby, S. Darling, A. Davis, L. Degenhardt, F. Dentener, D. C. Des Jarlais, K. Devries, M. Dherani, E. L. Ding, E. R. Dorsey, T. Driscoll, K. Edmond, S. E. Ali, R. E. Engell, P. J. Erwin, S. Fahimi, G. Falder, F. Farzadfar, A. Ferrari, M. M. Finucane, S. Flaxman, F. G. R. Fowkes, G. Freedman, M. K. Freeman, E. Gakidou, S. Ghosh, E. Giovannucci, G. Gmel, K. Graham, R. Grainger, B. Grant, D. Gunnell, H. R. Gutierrez, W. Hall, H. W. Hoek, A. Hogan, H. D. Hosgood, D. Hoy, H. Hu, B. J. Hubbell, S. J. Hutchings, S. E. Ibeanusi, G. L. Jacklyn, R. Jasrasaria, J. B. Jonas, H. Kan, J. A. Kanis, N. Kassebaum, N. Kawakami, Y.-H. Khang, S. Khatibzadeh, J.-P. Khoo, C. Kok, F. Laden, R. Lalloo, Q. Lan, T. Lathlean, J. L. Leasher, J. Leigh, Y. Li, J. K. Lin, S. E. Lipshultz, S. London, R. Lozano, Y. Lu, J. Mak, R. Malekzadeh, L. Mallinger, W. Marcenes, L. March, R. Marks, R. Martin, P. McGale, J. McGrath, S. Mehta, G. A. Mensah, T. R. Merriman, R. Micha, C. Michaud, V. Mishra, K. Mohd Hanafiah, A. A. Mokdad, L. Morawska, D. Mozaffarian, T. Murphy, M. Naghavi, B. Neal, P. K. Nelson, J. M. Nolla, R. Norman, C. Olives, S. B. Omer, J. Orchard, R. Osborne, B. Ostro, A. Page, K. D. Pandey, C. D. H. Parry, E. Passmore, J. Patra, N. Pearce, P. M. Pelizzari, M. Petzold, M. R. Phillips, D. Pope, C. A. Pope, J. Powles, M. Rao, H. Razavi, E. A. Rehfuess, J. T. Rehm, B. Ritz, F. P. Rivara, T. Roberts, C. Robinson, J. A. Rodriguez-Portales, I. Romieu, R. Room, L. C. Rosenfeld, A. Roy, L. Rushton, J. A. Salomon, U. Sampson, L. Sanchez-Riera, E. Sanman, A. Sapkota, S. Seedat, P. Shi, K. Shield, R. Shivakoti, G. M. Singh, D. A. Sleet, E. Smith, K. R. Smith, N. J. C. Stapelberg, K. Steenland, H. Stöckl, L. J. Stovner, K. Straif, L. Straney, G. D. Thurston, J. H. Tran, R. Van Dingenen, A. van Donkelaar, J. L. Veerman, L. Vijayakumar, R. Weintraub, M. M. Weissman, R. A. White, H. Whiteford, S. T. Wiersma, J. D. Wilkinson, H. C. Williams, W. Williams, N. Wilson, A. D. Woolf, P. Yip, J. M. Zielinski, A. D. Lopez, C. J. L. Murray, M. Ezzati, M. A. Al-Mazroa, and Z. A. Memish. A comparative risk assessment of burden of disease and injury attributable to 67 risk factors and risk factor clusters in 21 regions, 1990-2010: a systematic analysis for the Global Burden of Disease Study 2010. Lancet (London, England), 380(9859):2224–2260, 2012. ISSN 1474-547X. doi: 10.1016/S0140-6736(12)61766-8. URL http://www.ncbi.nlm.nih.gov/pubmed/23245609.

X. Liu, Y. Gao, J. Peng, Y. Xu, Y. Wang, N. Zhou, J. Xing, X. Luo, H. Jiang, and M. Zheng. TarPred: A web application for predicting therapeutic and side effect targets of chemical compounds. Bioinformatics, 31(12):2049–2051, jun 2015. ISSN 14602059. doi: 10.1093/bioinformatics/btv099. URL https://pubmed.ncbi.nlm.nih.gov/25686637/.

Y. C. Lo, S. E. Rensi, W. Torng, and R. B. Altman. Machine learning in chemoinformatics and drug discovery. Drug Discovery Today, 23(8):1538–1546, 2018. ISSN 18785832. doi: 10.1016/j.drudis.2018.05.010. URL https://doi.org/10.1016/j.drudis.2018.05.010.

C. J. Mattingly, G. T. Colby, J. N. Forrest, J. L. Boyer, A. A. P. A. Davis, C. J. Grondin, R. J. Johnson, D. Sciaky, R. McMorran, J. Wiegers, T. C. Wiegers, C. J. Mattingly, A. J. Williams, C. M. Grulke, J. Edwards, A. D. McEachran, K. Mansouri, N. C. Baker, G. Patlewicz, I. Shah, J. F. Wambaugh, R. S. Judson, A. M. Richard, F. Pedregosa, G. Varoquaux, A. Gramfort, V. Michel, B. Thirion, O. Grisel, M. Blondel, P. Prettenhofer, R. Weiss, V. Dubourg, J. Vanderplas, A. Passos, D. Cournapeau, M. Brucher, M. Perrot, É. Duchesnay, M. Abadi, P. Barham, J. S. Chen, Z. Chen, A. A. P. A. Davis, J. Dean, M. Devin, S. Ghemawat, G. Irving, M. Isard, M. Kudlur, J. Levenberg, R. Monga, S. Moore, D. G. Murray, B. Steiner, P. Tucker, V. Vasudevan, P. Warden, M. Wicke, Y. Yu, X. Zheng, Anaconda, Sylabs.io — Singularity, T. Liu, Y. Lin, X. Wen, R. N. Jorissen, M. K. Gilson, K. Preuer, P. Renz, T. Unterthiner, S. Hochreiter, G. Klambauer, A. Mayr, G. Klambauer, T. Unterthiner, M. Steijaert, J. K. Wegner, H. Ceulemans, D. A. Clevert, S. Hochreiter, S. Durinck, P. T. Spellman, E. Birney, W. Huber, M. Kuhn, D. Szklarczyk, A. Franceschini, M. Campillos, C. Von Mering, L. J. Jensen, A. Beyer, P. Bork, J. Nickel, B. O. Gohlke, J. Erehman, P. Banerjee, W. W. Rong, A. Goede, M. Dunkel, R. Preissner, S. Günther, J. Ahmed, B. Wittig, R. Preissner, X. Liu, Y. Gao, J. Peng, Y. Xu, Y. Wang, N. Zhou, J. Xing, X. Luo, H. Jiang, M. Zheng, H. Du, Y. Cai, H. Yang, H. Zhang, Y. Xue, G. Liu, Y. Tang, W. Li, E. Commission, R. Huang, M. Xia, D.-T. Nguyen, T. Zhao, S. Sakamuru, J. Zhao, S. Shahane, A. Rossoshek, A. Simeonov, G. Landrum, François Chollet, A. Author, Y. Another author, F. Authors, D. K. Barupal, O. Fiehn, S. M. Rappaport, D. K. Barupal, D. Wishart, P. Vineis, A. Scalbert, S. S. Lim, T. Vos, A. D. Flaxman, G. Danaei, K. Shibuya, H. Adair-Rohani, M. Amann, H. R. Anderson, K. G. Andrews, M. Aryee, C. Atkinson, L. J. Bacchus, A. N. Bahalim, K. Balakrishnan, J. Balmes, S. Barker-Collo, A. Baxter, M. L. Bell, J. D. Blore, F. Blyth, C. Bonner, G. Borges, R. Bourne, M. Boussinesq, M. Brauer, P. Brooks, N. G. Bruce, B. Brunekreef, C. Bryan-Hancock, C. Bucello, R. Buchbinder, F. Bull, R. T. Burnett, T. E. Byers, B. Calabria, J. Carapetis, E. Carnahan, Z. Chafe, F. Charlson, H. Chen, J. S. Chen, A. T.-A. Cheng, J. C. Child, A. Cohen, K. E. Colson, B. C. Cowie, S. Darby, S. Darling, A. A. P. A. Davis, L. Degenhardt, F. Dentener, D. C. Des Jarlais, K. Devries, M. Dherani, E. L. Ding, E. R. Dorsey, T. Driscoll, K. Edmond, S. E. Ali, R. E. Engell, P. J. Erwin, S. Fahimi, G. Falder, F. Farzadfar, A. Ferrari, M. M. Finucane, S. Flaxman, F. G. R. Fowkes, G. Freedman, M. K. Freeman, E. Gakidou, S. Ghosh, E. Giovannucci, G. Gmel, K. Graham, R. Grainger, B. Grant, D. Gunnell, H. R. Gutierrez, W. Hall, H. W. Hoek, A. Hogan, H. D. Hosgood, D. Hoy, H. Hu, B. J. Hubbell, S. J. Hutchings, S. E. Ibeanusi, G. L. Jacklyn, R. Jasrasaria, J. B. Jonas, H. Kan, J. A. Kanis, N. Kassebaum, N. Kawakami, Y.-H. Khang, S. Khatibzadeh, J.-P. Khoo, C. Kok, F. Laden, R. Lalloo, Q. Lan, T. Lathlean, J. L. Leasher, J. Leigh, Y. Li, J. K. Lin, S. E. Lipshultz, S. London, R. Lozano, Y. Lu, J. Mak, R. Malekzadeh, L. Mallinger, W. Marcenes, L. March, R. Marks, R. Martin, P. McGale, J. McGrath, S. Mehta, G. A. Mensah, T. R. Merriman, R. Micha, C. Michaud, V. Mishra, K. Mohd Hanafiah, A. A. Mokdad, L. Morawska, D. Mozaffarian, T. Murphy, M. Naghavi, B. Neal, P. K. Nelson, J. M. Nolla, R. Norman, C. Olives, S. B. Omer, J. Orchard, R. Osborne, B. Ostro, A. Page, K. D. Pandey, C. D. H. Parry, E. Passmore, J. Patra, N. Pearce, P. M. Pelizzari, M. Petzold, M. R. Phillips, D. Pope, C. A. Pope, J. Powles, M. Rao, H. Razavi, E. A. Rehfuess, J. T. Rehm, B. Ritz, F. P. Rivara, T. Roberts, C. Robinson, J. A. Rodriguez-Portales, I. Romieu, R. Room, L. C. Rosenfeld, A. Roy, L. Rushton, J. A. Salomon, U. Sampson, L. Sanchez-Riera, E. Sanman, A. Sapkota, S. Seedat, P. Shi, K. Shield, R. Shivakoti, G. M. Singh, D. A. Sleet, E. Smith, K. R. Smith, N. J. C. Stapelberg, K. Steenland, H. Stöckl, L. J. Stovner, K. Straif, L. Straney, G. D. Thurston, J. H. Tran, R. Van Dingenen, A. van Donkelaar, J. L. Veerman, L. Vijayakumar, R. Weintraub, M. M. Weissman, R. A. White, H. Whiteford, S. T. Wiersma, J. D. Wilkinson, H. C. Williams, W. Williams, N. Wilson, A. D. Woolf, P. Yip, J. M. Zielinski, A. D. Lopez, C. J. L. Murray, M. Ezzati, M. A. Al-Mazroa, and Z. A. Memish. Generating the blood exposome database using a comprehensive text mining and database fusion approach. Environmental Health Perspectives, 9(8):769–774, oct 2016. ISSN 13624962. doi: 10.1371/journal.pone.0154387. URL http://www.ncbi.nlm.nih.gov/pubmed/23245609.

A. Mayr, G. Klambauer, T. Unterthiner, and S. Hochreiter. DeepTox: Toxicity Prediction Using Deep Learning. Frontiers in Environmental Science, 3, 2015. doi: 10.3389/fenvs.2015.00080.

L. McInnes, J. Healy, N. Saul, and L. Großberger. UMAP: Uniform Manifold Approximation and Projection. Journal of Open Source Software, 3(29):861, 2018. ISSN 2475-9066. doi: 10.21105/joss.00861.

Norman Network. EMPODAT Database. https://www.norman-network.com/nds/empodat/chemicalStatistics.php, 2020.

F. Pedregosa Fabianpedregosa, V. Michel, O. Grisel Oliviergrisel, M. Blondel, P. Prettenhofer, R. Weiss, J. Vanderplas, D. Cournapeau, F. Pedregosa, G. Varoquaux, A. Gramfort, B. Thirion, O. Grisel, V. Dubourg, A. Passos, M. Brucher, M. Perrot andÉdouardand, A. Duchesnay, and F. Duchesnay EDOUARD-DUCHESNAY. Scikit-learn: Machine Learning in Python Gaël Varoquaux Bertrand Thirion Vincent Dubourg Alexandre Passos PEDREGOSA, VAROQUAUX, GRAMFORT ET AL. Matthieu Perrot. Technical report, 2011. URL http://scikit-learn.sourceforge.net.

R. Perkins, H. Fang, W. Tong, and W. J. Welsh. Quantitative structure-activity relationship methods: Perspectives on drug discovery and toxicology. Environmental Toxicology and Chemistry, 22(8):1666–1679, 2003. ISSN 07307268. doi: 10.1897/01-171.

L. Posthuma, J. van Gils, M. C. Zijp, D. van de Meent, and D. de Zwartd. Species sensitivity distributions for use in environmental protection, assessment, and management of aquatic ecosystems for 12 386 chemicals. Environmental Toxicology and Chemistry, 38(4):703–711, 2019. ISSN 15528618. doi: 10.1002/etc.4373.

L. Pu, M. Naderi, T. Liu, H.-C. Wu, S. Mukhopadhyay, and M. Brylinski. eToxPred: a machine learning-based approach to estimate the toxicity of drug candidates. BMC Pharmacology and Toxicology, 20(1), 2019. doi: 10.1186/s40360-018-0282-6.

A. B. Raies and V. B. Bajic. In silico toxicology: computational methods for the prediction of chemical toxicity. Wiley Interdisciplinary Reviews: Computational Molecular Science, 6(April):147–172, 2016. ISSN 17590884. doi: 10.1002/wcms.1240.

B. Ramsundar, P. Eastman, P. Walters, and V. Pande. Deep Learning for the Life Sciences. O’Reilly Media, Inc., 2019. ISBN 9781492039839. URL https://www.oreilly.com/library/view/deep-learning-for/9781492039822/.

S. M. Rappaport. Genetic Factors Are Not the Major Causes of Chronic Diseases. Plos One, 11(4):e0154387, 2016. ISSN 1932-6203. doi: 10.1371/journal.pone.0154387. URL http://dx.plos.org/10.1371/journal.pone.0154387.

L. Sun, H. Yang, Y. Cai, W. Li, G. Liu, and Y. Tang. In Silico Prediction of Endocrine Disrupting Chemicals Using Single-Label and Multilabel Models. Journal of Chemical Information and Modeling, 59, 2019. doi: 10.1021/acs.jcim.8b00551.

R. Thomas. The US Federal Tox21 Program: A strategic and operational plan for continued leadership. ALTEX, pages 163–168, 2018. ISSN 1868596X. doi: 10.14573/altex.1803011.

S. R. Vink, J. Mikkers, T. Bouwman, H. Marquart, and E. D. Kroese. Use of read-across and tiered exposure assessment in risk assessment under REACH-a case study on a phase-in substance. Regulatory toxicology and pharmacology: RTP, 58(1):64–71, 2010. ISSN 1096-0295. doi: 10.1016/j.yrtph.2010.04.004.

Z. Wang, G. W. Walker, D. C. Muir, and K. Nagatani-Yoshida. Toward a Global Understanding of Chemical Pollution: A First Comprehensive Analysis of National and Regional Chemical Inventories. Environmental Science and Technology, 54(5):2575–2584, mar 2020. ISSN 15205851. doi: 10.1021/ACS.EST.9B06379/SUPPL_FILE/ES9B06379_SI_001.PDF.

A. J. Williams, C. M. Grulke, J. Edwards, A. D. Mceachran, K. Mansouri, N. C. Baker, G. Patlewicz, I. Shah, J. F. Wambaugh, R. S. Judson, A. M. Richard, and W. A. Gov. The CompTox Chemistry Dashboard: a community data resource for environmental chemistry Open Access. Journal of Cheminformatics, 9:61, 2017. doi: 10.1186/s13321-017-0247-6.

Z. Wu, B. Ramsundar, E. N. Feinberg, J. Gomes, C. Geniesse, A. S. Pappu, K. Leswing, and V. Pande. MoleculeNet: A Benchmark for Molecular Machine Learning. Chemical Science, 9(2):513–530, mar 2017. URL http://arxiv.org/abs/1703.00564.

