## Supplement document for "AI for predicting chemical-effect associations at the chemical universe level – deepFPlearn"

This document contains detailed information about the used datasets, supplemental figures, and insights of the hyperparameter tuning of the autoencoder and feed-forward neural networks. General information is provided and clear explanations and observations of the figures comes with the figure captions. **keywords:**Deep learning, toxicology, binary fingerprint, autoencoder, molecular structures

### 1 Datasets

We provide the data that we used for training as CSV files:

1. *S_dataset.csv* contains all binary chemical – ED target associations that we used for training the specific deep autoencoder (AE) and the feed forward neural networks (FNNs). We have extracted the original data from the supplemental material of **?**.
2. *D_dataset.tsv* contains all InChI keys that we used to extract binary fingerprints for training the generic deep autoencoder (AE). These SMILES strings correspond to the chemical inventory of the CompTox Database (**?**, accessed on 2020/07/13).
3. *ER_predictions_CompTox.csv* and *ED_predictions_CompTox.csv* contain the prediction of all CompTox chemicals with the `deepFPlearn` models for estrogen receptor (ER) and general endocrine disruption (ED). We used the best models for each receptor from the 5fold cross validation study.

The FNN training data set originated from **?**. It contains chemical – target associations for six receptors that are involved in endocrine disruption: estrogen, androgen, glucocorticoid, thyroid receptors, peroxisome proliferator-activated receptor gamma, and aromatase, referred to as ER, AR, GR, TR, PPARg, and Aromatase. We summarized all individual targets to a more general endocrine disruption target - referred to as ED. See Fig. S1 for an overview of the distribution of labels in the training data that we used for FNN training. Based on the ratio between 1 and 0 class labels and the overall available amount of samples we decided to use AR, ER and ED for training the FNNs.

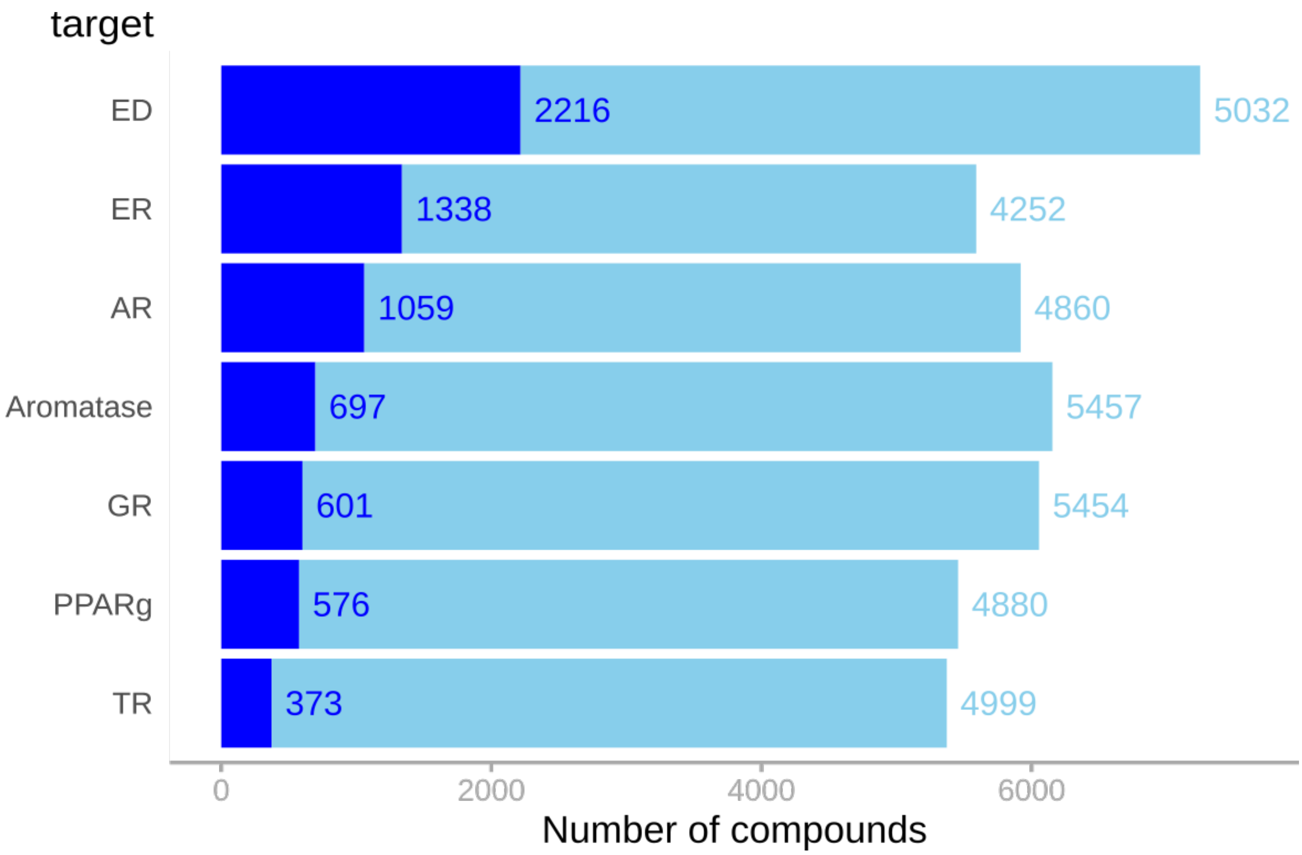


Fig. S1: Distribution of associations between compounds and endocrine disruption gene targets, stratified by type of association and separated by gene target. *Blue* colored bars count the compounds that are active regarding the gene target, *Lightblue* inactive compounds. This data has been extracted from **?** and extended by a general target for endocrine disruption (ED) which summarizes the individual targets with an OR operation.

#### 2 Model performances

We applied three different training setups for each target:

1. uncompressed - no feature compression, training of the FNN

2. compressed (*specific* AE) - feature compression based on the specific dataset from Sun et al. and training of t'he FNN

3. compressed (*generic* AE) - feature compression based on the chemical inventory from the CompTox Database and training of the FNN

For the FNNs, we employed 5-fold cross-validation to show that the selection of the train-test-split has no significant impact on the model performance. The standard deviation of the ROC-AUC values was ∼1%, see Fig. S2 for a comparison of the distribution and stdv of the ROC-AUC values for all 5 folds stratified by training setup and target.
This low variation among the folds allowed to use a single stratified train-test-split to finetune and train our models. We applied the trained models to the (unseen) validation data, and calculated confusion matrices, F1 scores, balanced accuracy and MCC values, and constructed the values for the respective precision-recall curves. See Fig. S3, Fig. S-4, and Fig. S5 for an overview across all selected targets (models).

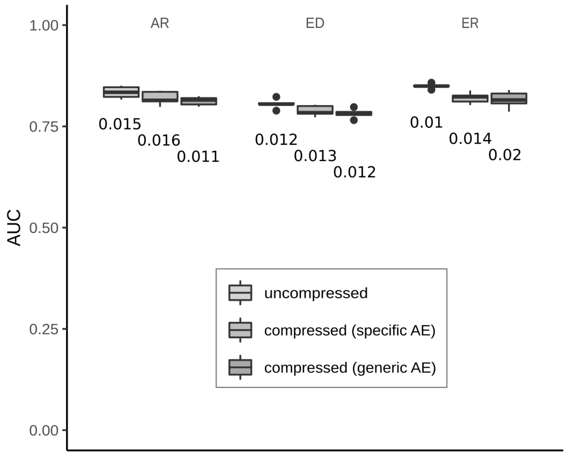


Fig. S2: Distribution of ROC-AUC values across the 5 folds of the cross-validation stratified by training setup and target/model. Shown values below the boxplots are the standard deviations of the ROC-AUC values.

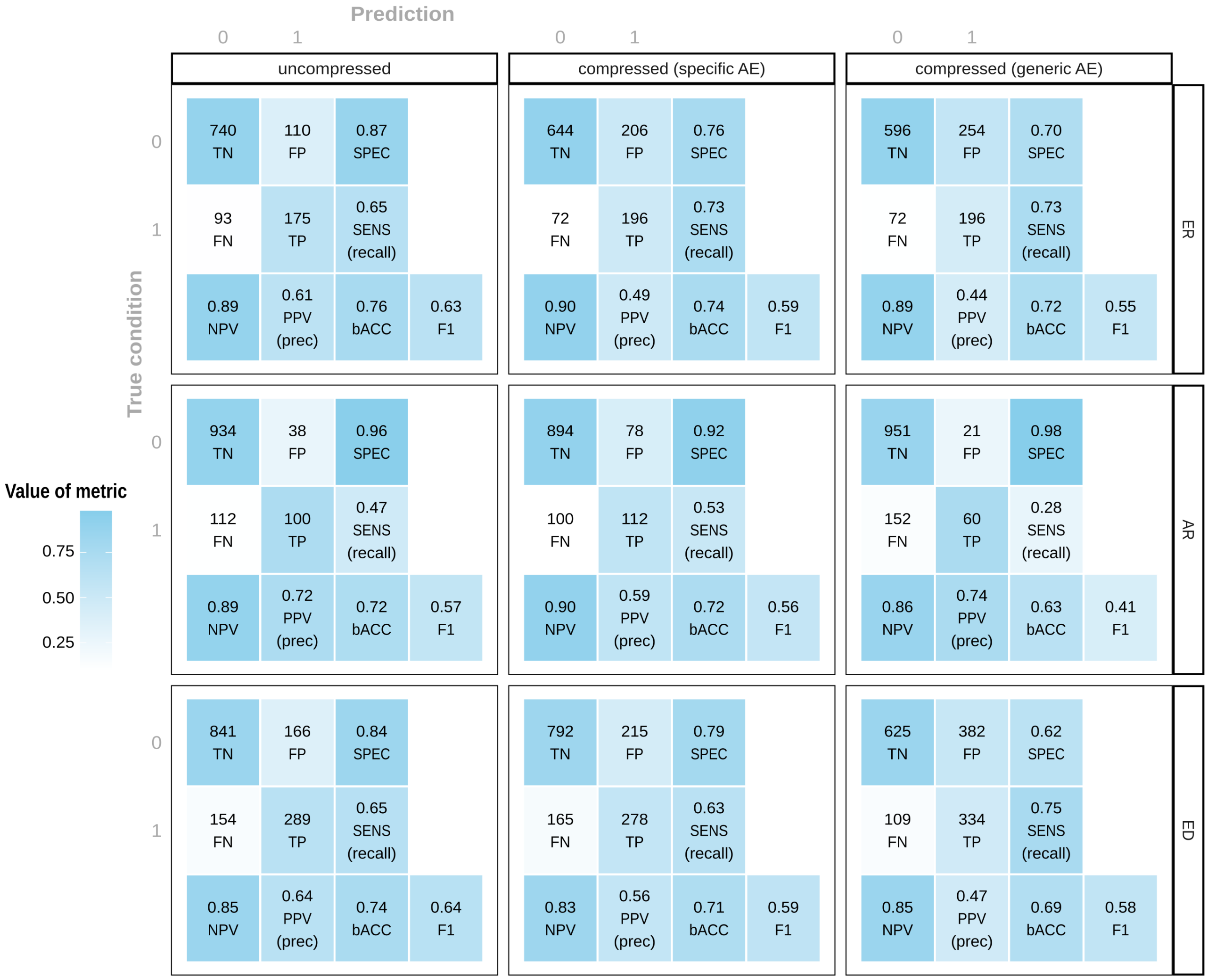


Fig. S3: CFMs calculated on respective validation data. Color scales between 0 and 1. Note that, TN, FP, FN, and TP are colored based on the underlying percentages of the printed count values, and that MCC scales from -1 to 1 but is colored based on its 0/1-scaled value.

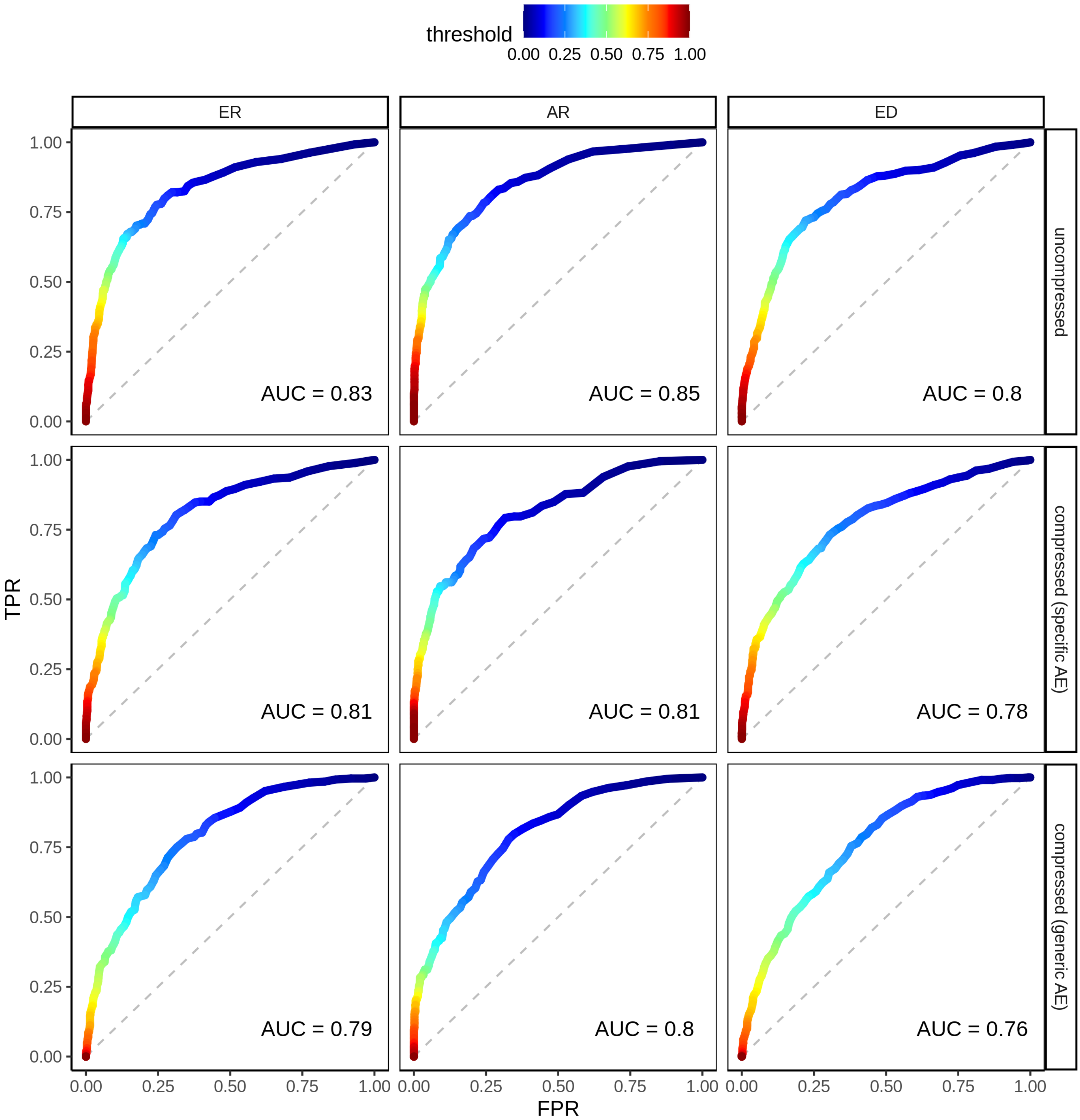


Fig. S4: ROC curves and ROC-AUC values stratified by the selected targets AR, ER, and ED, and by the degree of feature compression.

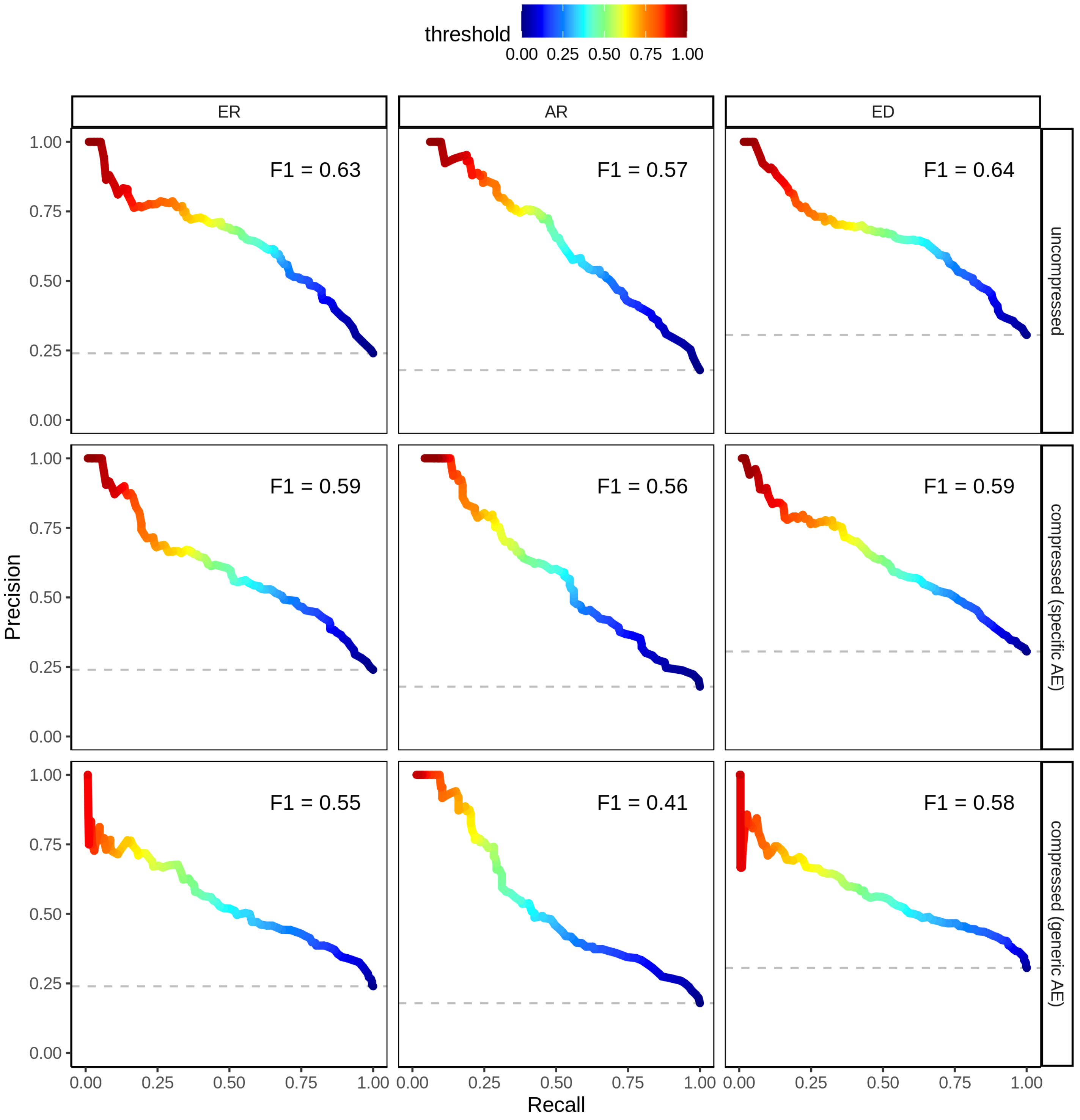


Fig. S5: Precision-recall curves and respective F1 scores stratified by the selected targets AR, ER, and ED, and by the degree of feature compression.

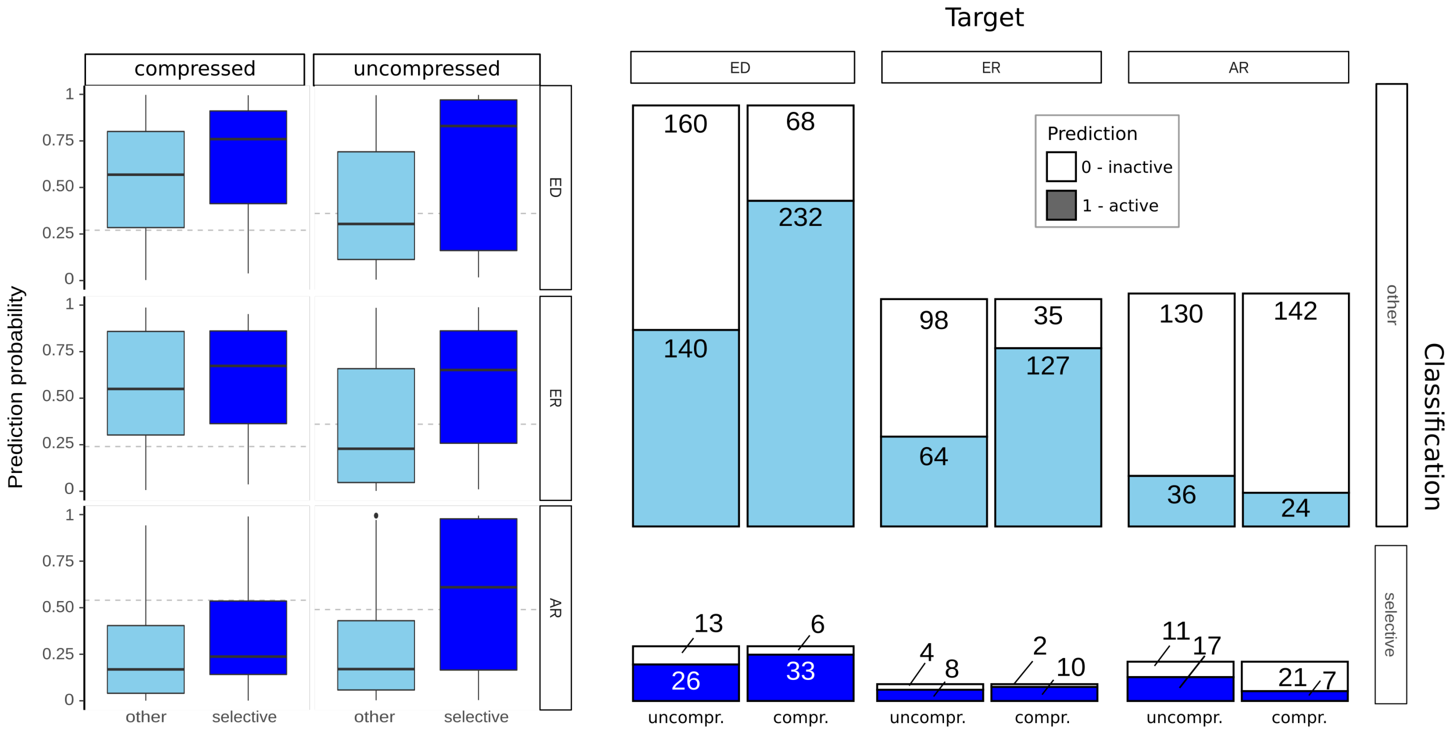


Fig. S6: Left: `deepFPlearn` prediction probability of the predicted association of the compounds that have been experimentally verified for receptor activity by ?. Right: Comparison of the counts of predicted 1 (active) and 0 (inactive) labels for the same compounds. The color refers to the differentiation into selective and other binders by ?.

### 3 Feature compression preserves structural information

We investigated the clustering of the input feature vector and compared it to a clustering of the compressed feature vector. When coloring compounds both from the uncompressed and compressed space with labels that were calculated on the uncompressed feature space, it is clearly visible that a similar cluster association appears in the UMAP. This strongly indicates that relevant structural information is preserved during feature compression. Fig. **??** shows the results of the clustering for a different number of clusters ($k \in [2, 3, 5, 7]$) visualized as a UMAP embedding.

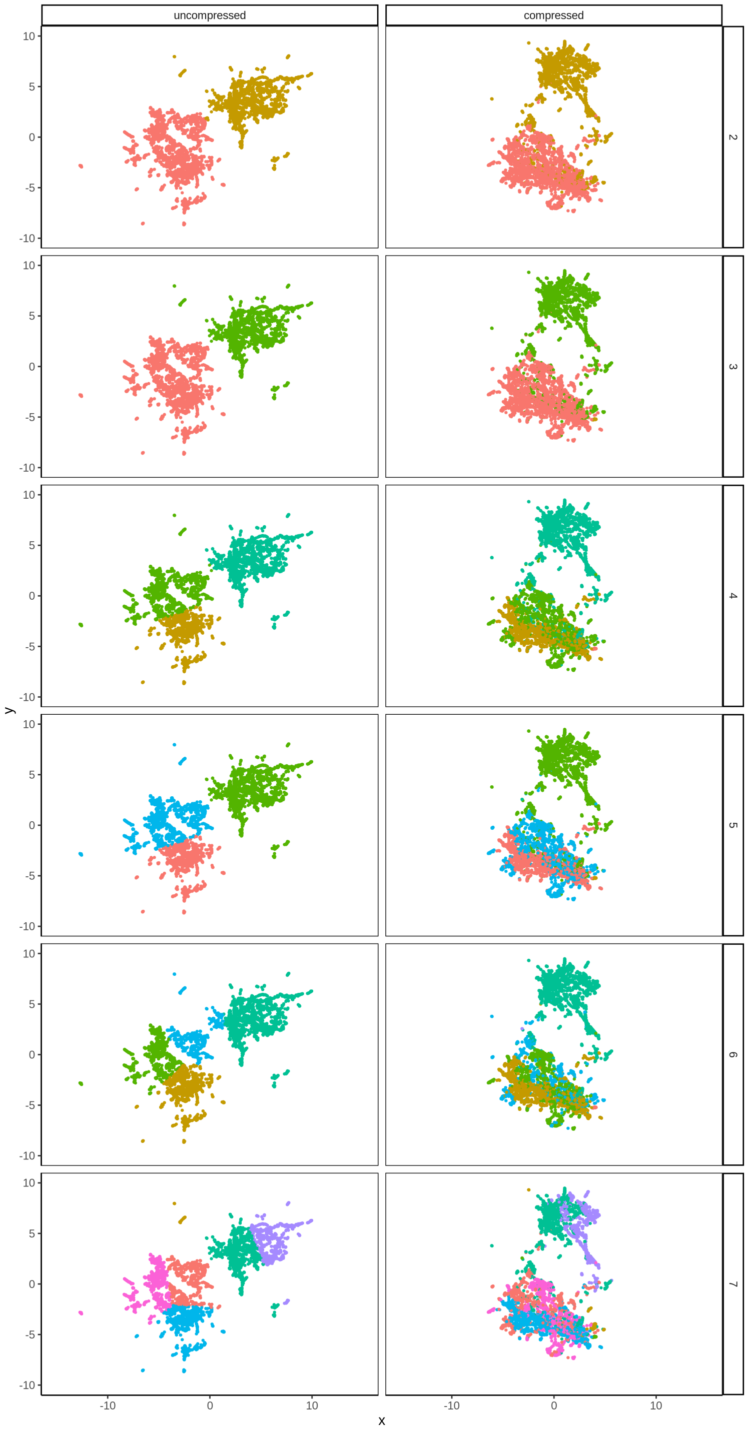


Fig. S7: Feature compression preserves relevant structural information. UMAP visualizations of uncompressed and compressed representations of all compounds from $S$ dataset; color indicates cluster assignment of a $k$-means clustering with $k \in [2, 3, 5, 7]$ on the uncompressed features. UMAP visualization of uncompressed and compressed features of all the compounds from the $S$ dataset.

### 4 Hyperparameter tuning

In the following, we will give insight into the process of hyperparameter-tuning for both the auto-encoders and the feed-forward networks used in the manuscript. When the general architectures of the used networks were established, parameters like batch sizes, learning rates, learning rate decays, dropout probabilities or activation functions that influence the performance needed to be optimized.

In general, we followed the strategy of making random sweeps through a wide range of parameters to filter out candidates that did not converge well. After the initial sweep, we fixed certain hyperparameters like using the Adam optimizer in favour of SGD. For others, the sensible range was narrowed down drastically. At this point a Bayesian search through the hyperparameter space was employed to find optimal settings.

Overall, we trained over 3000 networks to track and optimize performance using *Weights and Biases* (https://wandb.ai).

#### Autoencoder (AE) training

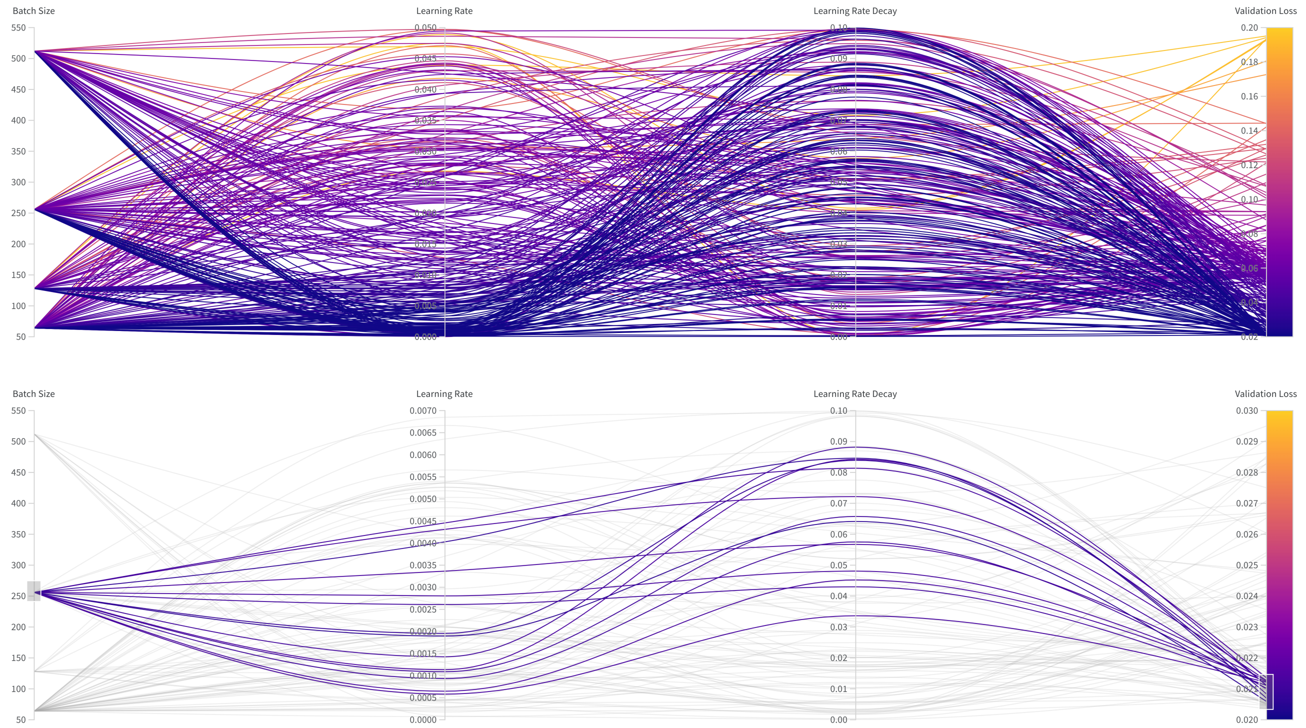


Fig. S8: Sweep through hyperparameter values for the auto-encoder (AE). Top panel shows a wide view of parameters tested, while the bottom panel shows a filtered view indicating the parameter ranges for the best-performing runs.

After the initial sweep, SELU activation was used in favor of RELU since we observed a better convergence of training runs. Therefore, the main training parameters to optimize for were the batch size, the learning rate and its decay.

Fig. **??** shows how different values for these hyperparameters influence the validation loss of the overall training. The training was allowed to run for 3000 epochs but could end early when the validation loss stopped to improve. From the upper panel of this figure it is apparent that smaller learning rates will result in better performance, while the correlation of the learning rate decay with the validation loss was low. After applying a filter for a particular batch size and the best performing runs, which is shown in the bottom panel of Fig. **??**, we retrieved a much better impression of which ranges of learning rate and its decay work best for a particular batch-size.

Fig. **??** shows validation metrics for 10 runs of the narrow selection shown in the bottom of Fig. **??**. As can be seen, all of these runs perform well but a good choice should have a smooth convergence rather than training as fast as possible.

For final hyperparameter values used in the manuscript, we tried to balance smooth convergence with high performance in the validation metrics.

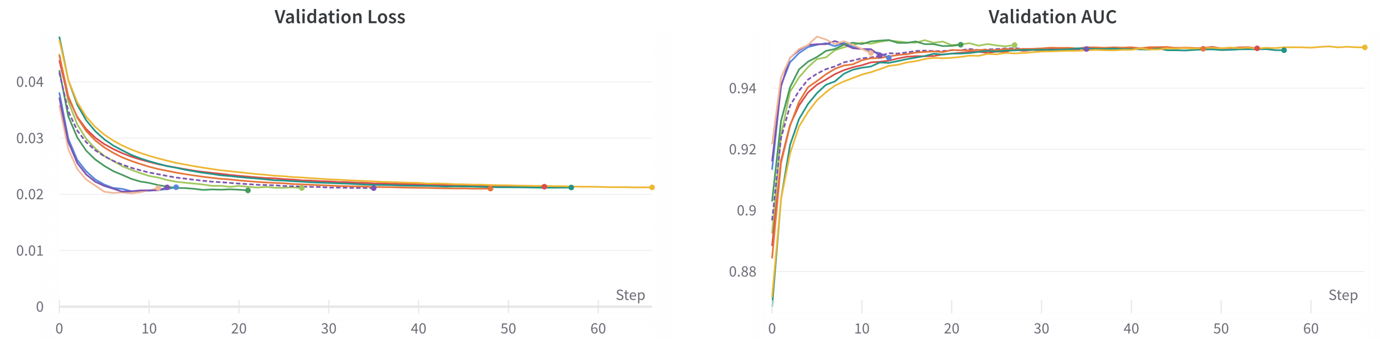


Fig. S9: Validation loss and AUC for 10 of the best performing runs shown in the bottom panel of Fig. ??.

#### Feedforward network (FNN) training

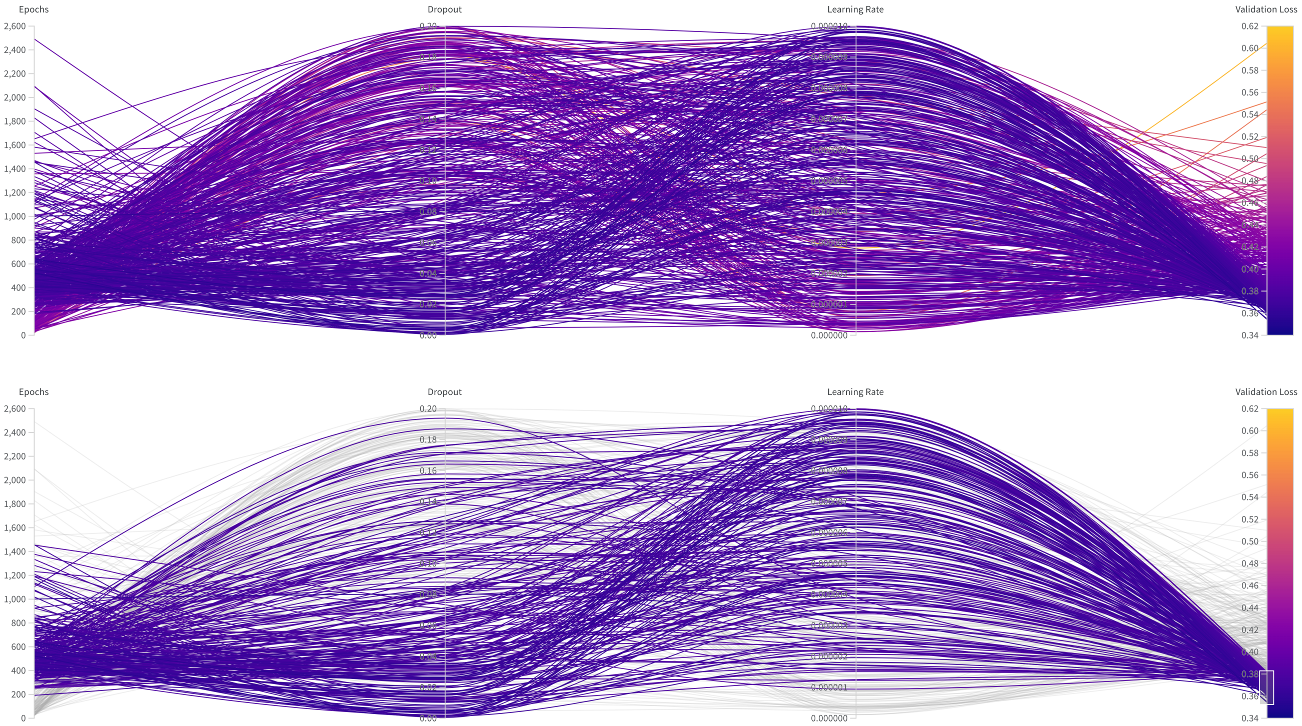


Fig. S10: Sweep through hyperparameter values for the feed-forward networks. Top panel shows a view of wide view of of parameters tested, while the bottom panel shows a filtered view indicating the parameter ranges for the best-performing runs.

For the feedforward networks (FNN) we followed a similar approach as for the AEs. First, an initial random sweep to narrow down possible candidates that performed better than others before running a narrow sweep to find good hyperparameter values. In the case of the FNNs we fixed the batch-size to 128, used SELU activation and used the default setting for the learning rate decay since it did not have a strong influence as it had when testing the AE. The two major parameters to sweep were the learning rate and the dropout probability.

Since we have seen in the AE section that the number of epochs trained influences the smoothness of the convergence, Fig. ?? also shows for how long the training ran. The number of epochs was rather a result of the early stopping and not a hyperparameter that we chose. However, we used it to discard candidates that trained unnecessary long or too fast to show a steady convergence.

Fig. ?? shows that good convergence was achieved for a wide range of hyperparameters, but using a lower dropout probability and a learning rate in the upper values of the shown range is more likely to converge to a low validation loss within 300 to 700 epochs.

Fig. ?? shows two selected runs from the filtered selection in the bottom panel of Fig. ??. It clearly indicates that both the training and the validation metrics have a smooth and steady convergence.

#### The final selection of parameter values

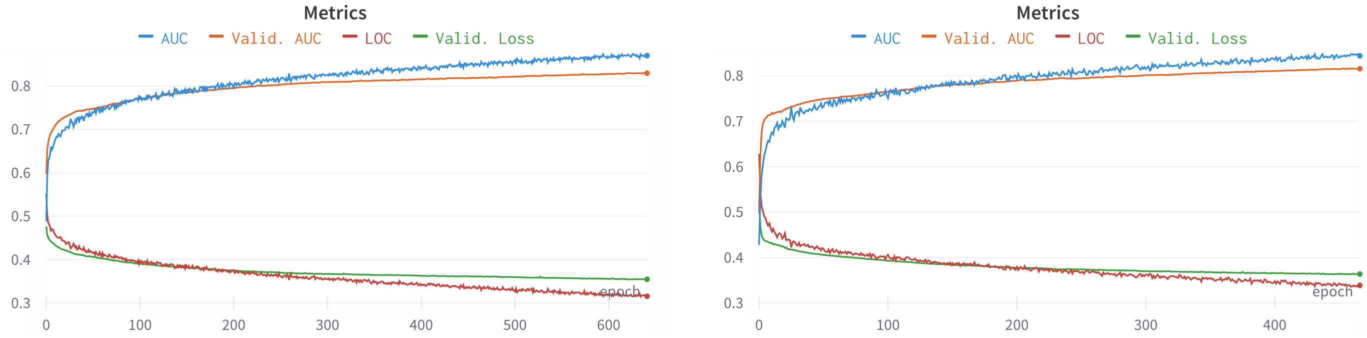


Fig. S11: Two selected runs for the FNNs showing the performance using different hyperparameters selected from promising candidates shown in Fig. **??**.

| **Autoencoder** | | | **Feed forward neural network** | | |
|---|---|---|---|---|---|
| | specific | generic | | uncompressed | compresssed |
| Batch size | 256 | 256 | Batch size | 128 | 128 |
| Learning rate | 0.0007 | 0.0011 | Learning rate | 0.0000022 | 0.0000078 |
| Learning rate decay | 0.0023 | 0.0600 | Dropout | 0.0107 | 0.0238 |

Table 1: Overview of all parameter settings for training the AEs and the FNNs that result from our hyperparameter tuning and were selected for the final training of all our models.

### References


B. I. Escher, L. Henneberger, M. König, R. Schlichting, and F. C. Fischer. Cytotoxicity burst? Differentiating specific from nonspecific effects in tox21 in vitro reporter gene assays. *Environmental Health Perspectives*, 128 (7):1–10, jul 2020. ISSN 15529924. doi: 10.1289/EHP6664. URL `https://ehp.niehs.nih.gov/doi/10.1289/EHP6664`.

L. Sun, H. Yang, Y. Cai, W. Li, G. Liu, and Y. Tang. In Silico Prediction of Endocrine Disrupting Chemicals Using Single-Label and Multilabel Models. *Journal of Chemical Information and Modeling*, 59, 2019. doi: 10.1021/acs.jcim.8b00551.

A. J. Williams, C. M. Grulke, J. Edwards, A. D. Mceachran, K. Mansouri, N. C. Baker, G. Patlewicz, I. Shah, J. F. Wambaugh, R. S. Judson, A. M. Richard, and W. A. Gov. The CompTox Chemistry Dashboard: a community data resource for environmental chemistry Open Access. *Journal of Cheminformatics*, 9:61, 2017. doi: 10.1186/s13321-017-0247-6.
